# Stochastic satisficing account of confidence in uncertain value-based decisions

**DOI:** 10.1101/107532

**Authors:** Uri Hertz, Bahador Bahrami, Mehdi Keramati

## Abstract

Every day we make choices under uncertainty; choosing what route to work or which queue in a supermarket to take, for example. It is unclear how outcome variance, e.g. uncertainty about waiting time in a queue, affects decisions and confidence when outcome is stochastic and continuous. How does one evaluate and choose between an option with unreliable but high expected reward, and an option with more certain but lower expected reward? Here we used an experimental design where two choices’ payoffs took continuous values, to examine the effect of outcome variance on decision and confidence. We found that our participants’ probability of choosing the good (high expected reward) option decreased when the good or the bad options’ payoffs were more variable. Their confidence ratings were affected by outcome variability, but only when choosing the good option. Unlike perceptual detection tasks, confidence ratings correlated only weakly with decisions’ time, but correlated with the consistency of trial-by-trial choices. Inspired by the satisficing heuristic, we propose a “stochastic satisficing” (SSAT) model for evaluating options with continuous uncertain outcomes. In this model, options are evaluated by their probability of exceeding an acceptability threshold, and confidence reports scale with the chosen option’s thus-defined satisficing probability. Participants’ decisions were best explained by an expected reward model, while the SSAT model provided the best prediction of decision confidence. We further tested and verified the predictions of this model in a second experiment. Our model and experimental results generalize the models of metacognition from perceptual detection tasks to continuous-value based decisions. Finally, we discuss how the stochastic satisficing account of decision confidence serves psychological and social purposes associated with the evaluation, communication and justification of decision-making.

**Author Summary:** Every day we make several choices under uncertainty, like choosing a queue in a supermarket. However, the computational mechanisms underlying such decisions remain unknown. For example, how does one choose between an option with unreliable high expected reward, like the volatile express queue, and an option with more certain but lower expected reward in the standard queue? Inspired by bounded rationality and the notion of ‘satisficing’, i.e. settling for a good enough option, we propose that such decisions are made by comparing the likelihood of different actions to surpass an acceptability threshold. When facing uncertain decisions, our participants’ confidence ratings were not consistent with the expected outcome’s rewards, but instead followed the satisficing heuristic proposed here. Using an acceptability threshold may be especially useful when evaluating and justifying decisions under uncertainty.

## Introduction

Every morning most people have to pick a route to work. While the shortest route may be consistently busy, others may be more variable, changing from day to day. The choice of which route to take impacts the commuting time and is ridden with uncertainty. Decision making under uncertainty has been studied extensively using scenarios with uncertain rewards [1–3]. In such scenarios, participants choose between multiple lotteries where each lottery can lead to one of the two (or several) consequences with different probabilities. Standard models like expected utility theory [4,5] and prospect theory [6] suggest parsimonious formulations for how the statistics of such binomial (or multinomial) distributions of outcomes determine the value (otherwise known as utility) of a lottery. These models, for example, explain the fact that in certain ranges people prefer a small certain reward to bigger more uncertain ones [7,8].

However, the commuting problem described here highlights the pervasive but much less studied relevance of outcome variance to decisions with continuous (rather than binary) outcomes. It is not straightforward how one’s choice and evaluation of the route could be decided using the heuristics applicable to binary win/lose outcomes.

Early studies of bounded rationality [9–11] introduced the concept of satisficing according to which, individuals replace the computationally expensive question of “which is my best choice?” with the simpler and most-of-the-times adequately beneficial question “which option is good enough?”. More precisely, instead of finding the best solution, decision makers settle for an option that satisfices an acceptability threshold [9]. In the case of commuting, such acceptability threshold could be “the latest time one affords to arrive at work". A generalization of the satisficing theory to decision-making under uncertainty suggests that when outcomes are variable, one could evaluate - with reasonably simple and general assumptions about the probability distributions of outcomes - the available options’ *probability* of exceeding an acceptability threshold [12,13]. Our commuter’s stochastic satisficing heuristic could then be expressed as “which route is *more likely* to get me to work before X o’clock?”

The effect of uncertainty on confidence report is commonly studied in perceptual detection tasks where one has to detect a world state from noisy stimuli (e.g. dots moving to the left or right) [14–20]. Sanders and colleagues (2016) argued that confidence report in perceptual decisions relates to the Bayesian formulation of confidence used in hypothesis testing. In this view, subjective confidence conveys the posterior probability that an uncertain choice is correct, given the agent’s prior knowledge and noisy input information. Generalizing this scheme to value-based contexts, our probabilistic satisficing heuristic is naturally fit to account for the computational underpinnings of choice confidence and draws strong predictions about how confidence would vary with outcome variance. In fact, if choices were made by the probabilistic satisficing heuristic described above, confidence in those choices would be directly proportional to the probability that the chosen option exceeded the acceptability threshold. A choice whose probability of exceeding the acceptability threshold is higher should be made more confidently than another that barely passes the criterion, even if they have equal expected values.

Here we asked if, and how, human decision makers learn and factor outcome variance in their evaluation of choices between options with independent continuous returns. We hypothesised that decision makers use a stochastic satisficing heuristic to evaluate their choices and that their confidence conveys the estimated probability of the chosen option’s value to exceed the satisficing criterion. In two experiments, we used two-armed bandit tasks in which the expected values and variances associated with outcomes of each arm were systematically manipulated. We tested the stochastic satisficing model against a reward maximizing model [4,5,13,21] and an expected utility model [22–24] that propose alternative ways of computing choice and confidence as a function of the estimated statistics of options’ returns.

## Results

Participants performed a two-armed bandit task online where rewards were hidden behind two doors (Fig 1A) and the reward magnitudes followed different probability distributions (Fig 1B). On each trial, the participant decided which door to open, and expressed their choice confidence using a combined choice-confidence scale. Choosing the left side of the scale indicated choice of the left door and distance from the midline (ranged between 1: uncertain, to 6: certain) indicated the choice confidence. After the decision, the reward behind the chosen door was revealed and a new trial started. Each experimental condition was devised for a whole block of consecutive trials during which the parameters (mean and variance) governing the reward distribution for each door were held constant. Each block lasted between 27 and 35 trials (drawn from a uniform distribution). Transition from one block to the next happened seamlessly and participants were not informed about the onset of a new block.

**Fig 1:**
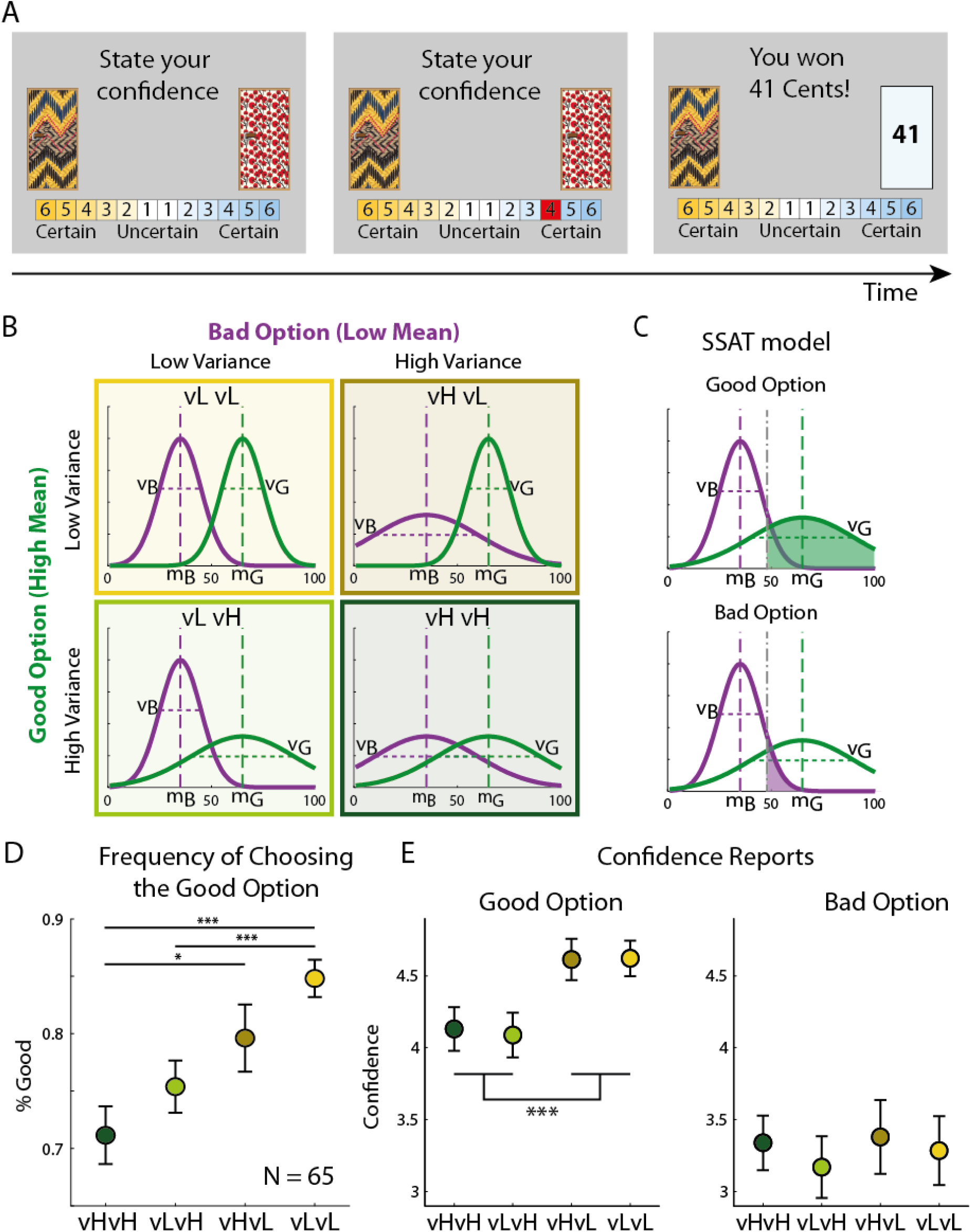
Design and results of experiment 1. (A) On each trial participants had to choose between two doors, using a confidence scale. The choice was determined by the side of the scale used by the participant. Upon decision, the chosen door was opened and the reward was revealed. See a working demo at http://urihertz.net/BanditConfDemo/ (B) Four different experimental conditions were embedded in a continuous two-armed bandit task. In each condition, one door had a low expected reward (Bad option) and the other had a high expected reward (Good option). Expected rewards (mB and mG) were constant across conditions. The variances of the two distributions, however, changed across conditions and were either high or low, resulting in a 2 × 2 design (VB (Low/High) x VG (Low/High)). Each condition lasted between 27 to 35 consecutive trials. (C) Stochastic satisficing model suggests that decisions are evaluated based on the probability of each door’s outcome exceeding an acceptability threshold (grey dot-dashed line). This probability (area under the curve) is higher for the door with the high mean expected reward (top) than for the door with the low mean (bottom). (D) Participants’ frequency of choosing the good option in each experimental condition, averaged across trials 10 to 25. (E) Participants’ confidence reports when choosing the good (middle panel) or bad (right panel) option. Reports were averaged between trials 10 to 25 of each experimental block. When choosing the good option, confidence ratings were higher when variance of the good option was low, regardless of the variance of the bad option. Confidence reports were not significantly different across conditions when choosing the bad option. Error bars represent SEM (* p<0.05, *** p<0.0005)

### Experiment 1

Within each condition, the rewards behind the doors were drawn from Gaussian distributions, one with a higher mean (65, i.e. the “good” option) than the other door (35, i.e. the “bad” option). The variances of the bad and good options could independently be high (H=25^2^=625) or low (L=102=100), resulting in a 2 × 2 design comprising four experimental conditions: ‘vH-vH’, ‘vL-vH’, ‘vH-vL’ and ‘vL-vL’ (Fig. 1B). In this notation, the first Capital letter indicates the variance of the bad (low expected value) option, and the second letter indicates the variance of the good (high expected value) options. Participants’ trial-by-trial probability of choosing the good option, in each condition, started at chance level and increased with learning until it reached a stable level after about 10 trials. To assess the level of performance after learning, we averaged the probability of choosing the good option between trials 10 to 25 in each experimental condition. Probability of choosing the good option was highest in the ‘vL-vL’ and lowest during the ‘vH-vH’ condition (Fig 1D). A repeated measure ANOVA test with the variances of the good and bad options as within-subject factors was used to evaluate this pattern. The effects of both variance factors were significant (variance of good option: F(1,194)=22.24, p=0.00001, variance of bad option: F(1,194)=5.2, p=0.026). This result indicated an asymmetric effect of outcomes’ payoff variances on choice: increased variance of the good option reduced the probability of choosing the good option, whereas increased variance of the bad option increased the probability of choosing the bad option. This variance-dependent choice pattern demonstrates that decision-making depended not only on the expected rewards, but also on their variances.

To examine the pattern of confidence reports, we calculated the average confidence reported on trials 10 to 25 in each condition (Fig 1E). Using a repeated measures ANOVA we found that the main effect of the good, but not the bad, option’s reward was significant when choosing the good option (good option variance: F(1,194)=33.32, p<0.00001, bad option variance effect: F(1, 194)=0.02, p=0.89). When choosing the bad option, confidence ratings were generally lower (paired t-test t(64)=8.3, p=10-^12^) and were not significantly different across experimental conditions. Therefore, variance affected confidence reports, but only when choosing the good option.

Several studies in perceptual decision making reported a strong relationship between decision confidence and reaction time [16,19]. We tested the correlation between each participant’s trial-by-trial reaction times and their confidence ratings (Fig 2A). Consistent with previous results, we found that the participants’ correlation coefficients tended to be below 0, as fast responses were linked to higher confidence (t-test, p=0.0015 in experiment 1). However, these were not as strongly linked across the population, with average correlation coefficient of R = −0.05, below the critical value of R(240) = 0.13 for significance of 0.05.

**Fig 2:**
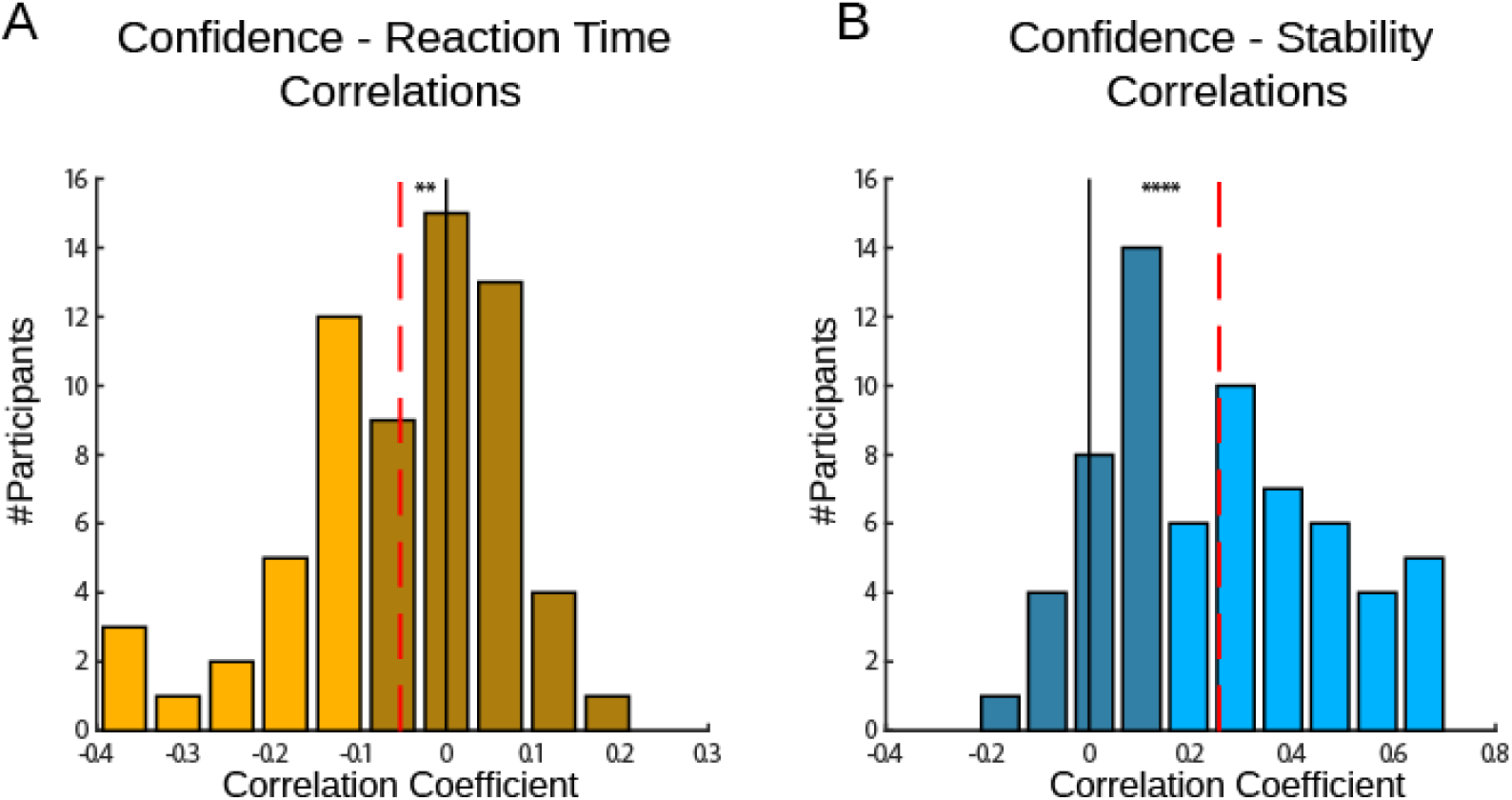
Correlations between confidence, reaction time, and stability in Experiment 1. (A) We correlated each participant’s reaction times with confidence ratings. We found that the participants’ correlation coefficients tended to be below 0, as fast responses were associated with higher confidence. However, these were not as strongly linked across the population, with average correlation of R=−0.05 (dashed line). (B) We examined the relations between confidence and choice stability. We found that the participants’ correlation coefficients were highly significant in the individual level and in the group level (average R = 0.26, dashed line). Dashed red lines indicate the average correlation coefficient. Dark colours indicate below significance correlation (Critical value of R(240) = 0.13, p = 0.05). ** p < 0.005, **** p < 0.00005.

We examined the relations between confidence and another behavioural measure – choice stability. We summed the number of choice switches in five trials sliding window, and subtract it from five to define the trial-by-trial choice stability. This measure ranged between 5, when no switches were made and the same option was chosen on all five trials, and 1 when the participant switched between every trial in the five trials window. We correlated each participant’s trial-by-trial stability measure with confidence reports (Fig 2B). We found that the participants’ correlation coefficients were highly significant in the individual level and in the group level (average R = 0.26 in experiment 1, p = 10-^13^).

### Fitting models to choice

To examine the use of a probabilistic satisficing heuristic and acceptability threshold in decisions under uncertainty we devised a set of models competing models (See Methods). We started with a model aimed at maximizing rewards (‘Reward’ model, hereafter) that tracks the expected reward from each door on every time-step [13,21,25]. Choice is then made according to the expected reward of each option [4]. We also tested an expected utility model (‘Utility’ model) which penalized options for their payoff’s variance, according to the participant’s risk-averse attitude [4,22,23] (In the supplementary materials we describe the performance of another variant of expected utility model, using power utility function instead of exponential utility function, Fig S6).

We added two other models, ‘Reward–T’ and ‘Utility-T’, to our set of competing models by formalizing the use of acceptability threshold and adding a free parameter ‘threshold’ to the ‘Reward’ and ‘Utility’ models. In these models, on each trial, the unchosen option drifted towards the value of this parameter. This ‘threshold’ therefore represented the participant’s expectations of outcome in the game. When this threshold was very high, the participant would assume that the unchosen option drifted toward this high threshold, making him likely to switch options often. A participant with a low threshold, however, might stick to an option even if it yielded low reward, as since the unchosen option would drift towards an even lower threshold.

We formalized the probabilistic satisficing heuristic in a stochastic satisficing (‘SSAT’) model [13] in which the mean and variance of rewards obtained by each of the two doors are tracked in a trial-by-trial manner (see Methods). In this model, decision was made by comparing the probability of each option yielding a reward above an acceptability threshold, i.e. being good enough. In fact, given the estimated probability distribution over the rewarding outcome of a choice, the model computed the total mass under this distribution that is above the acceptability threshold (Fig 1C). This cumulative quantity was then used to determine the probability of choosing that option. Such mechanism can capture the asymmetric effect of payoff variance on choice, as the good option (i.e. higher than threshold) becomes less likely to exceed the acceptability threshold as its variance increases, while the bad option (below threshold) becomes more likely to exceed the threshold as its variance increases. Upon making the choice and receiving reward feedback from the environment, the model updates the distribution over the value of the chosen action. We also examined a drift version of the SSAT model in which the value of the unchosen action drifts toward the acceptability threshold (‘SSAT-T’), similar to the drifting mechanism described above for ‘Reward-T’ and ‘Utility-T’ models.

In the light of previous theoretical studies that examined the optimality of the satisficing, maximizing and risk aversive models in accruing rewards [13,23,26], we examined our models’ performance in the experimental design. We also examined the rewards accrued by a model with full knowledge of the reward distribution (‘Omniscient’), and the actual amount of reward accrued by our participants. For each model we identified the set of parameters that maximized the model’s accumulated reward under the reward distributions in Experiment 1. We used these parameters to simulate each model, and compared the amount of reward accrued by each model (Fig S1). In accordance with the theoretical optimality analysis [13,23], we found that all three models performed similarly, and accrued similar amount of reward. When the drifting mechanism was added (drift of the unchosen option towards the acceptability threshold) performance of all models decreased. None of the models accrued as much reward as the ‘omniscient’ model, as all of them had to learn the statistics of the changing environment. In addition, all models performed much better than our participants, indicating that participants’ behaviour was noisy, falling short of the optimal strategy.

Another line of analysis was carried out to elucidate the differences between the models and how they operate in our experimental design. We examined the differences in values assigned to each option in a steady state (i.e. after learning the reward distributions) calculated by each model in the four conditions of our experimental design (Fig S2). We found that all models assigned higher values to the high mean reward option than to the low mean reward option in almost all the cases and conditions. An optimal (greedy) decision maker would therefore be able to accumulate similar amount of rewards using either model. However, the value differences assigned by each model were different in magnitude, if not in direction. This means that a noisy/exploratory decision maker (e.g. softmax) may be more likely to choose the low mean reward option in one condition compared with other conditions. Taken together, the two analyses demonstrate that all models have a similar potential for collecting reward under optimal condition and decision policy. However, our second analysis provided a prediction of the pattern of suboptimal choice when decisions are noisy.

We fitted all models to the choices made by the participants (240 trials per participant, model fitting was done for each participant independently) using Monte-Carlo-Markov-Chain (MCMC) procedure [27]. After correction for the number of parameters using Watanabe-Akaike information criterion (WAIC) [28], we compared posterior likelihood estimates obtained for each participant, for each model (Table 1). The models using drifting thresholds all performed better (lower WAIC score) than those not using it, and the ‘Reward-T’ model gave the best fit to the choice data (paired t-test vs. Reward p = 0.00003, vs. Utility p = 0.00001, vs. Utility-T p = 0.006, vs. SSAT p = 0.0002, vs. SSAT-T p = 0.0002, Fig 3A). We examined how many participants’ choices were best explained by each of our six models and found that while some participants’ behaviour was better explained by one of the other models, most of the participants’ choices were best explained by models that did not track reward variance, in line with the model comparison results (Fig S3).

**Table 1:** Models performance in Experiment 1.

| Model | Sum WAIC | Mean ± STD WAIC | Mean ± STD Confidence $R^2$ |
| --- | --- | --- | --- |
| Reward | 14,273 | 226.55 ± 67.67 | 0.21 ± 0.22 |
| Utility | 14,320 | 227.29 ± 67.48 | 0.18 ± 0.18 |
| SSAT | 14,047 | 222.96 ± 67.96 | 0.21 ± 0.22 |
| Reward-T | **13,518** | 214.57 ± 68.79 | 0.21 ± 0.21 |
| Utility-T | 13,813 | 219.25 ± 64.38 | 0.17 ± 0.16 |
| SSAT-T | 13,695 | 217.38 ± 67.61 | **0.24 ± 0.21** |

**Fig 3:**
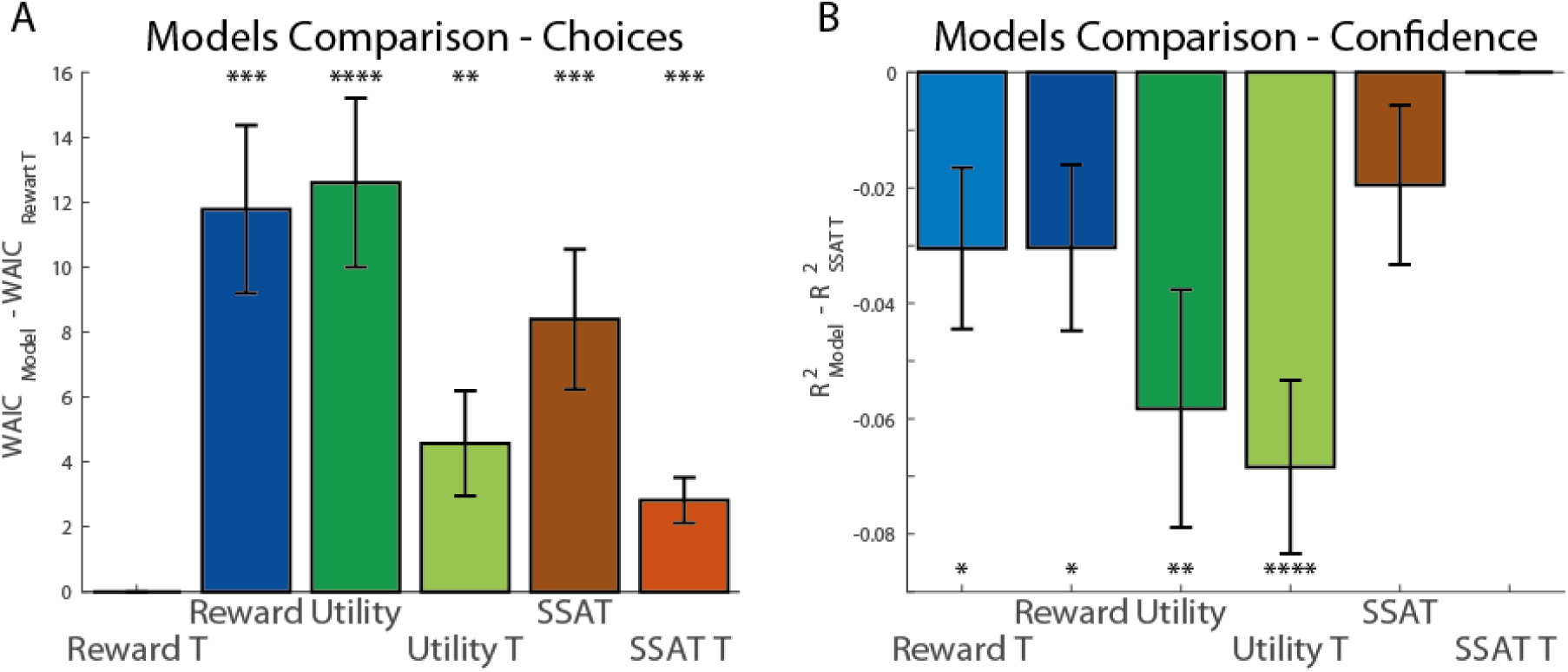
Models comparison in experiment 1. (A) We compared the ‘Reward T’ model to all the other models by examining the paired differences in WAIC scores across models and participants. The graph presents the differences of each model’s WAIC from the ‘Reward T’ model. The ‘Reward T’ model performed significantly better than all other models in explaining participants’ choices. (B) We compared the ‘SSAT T’ model to all other models by examining the paired differences in R^2^ scores across models and participants. The ‘SSAT T’ model gave a significantly better prediction of confidence reports than all other models except the ‘SSAT’ model. Error bars represent SEM. * p<0.05, ** p<0.005, *** p<0.0005, **** p<0.00005.

Examining the individual parameters fitted by the models we observed a high correspondence between the ‘Reward-T’ and ‘SSAT-T’ models for both the parameters estimated for acceptability threshold and the parameters estimated for learning rate (Table 2). This correspondence was captured by high correlation between the individual threshold parameters (R^2^=0.89) and learning-rate parameters (R^2^=0.86) (Fig S4). Such similarity was not found between the ‘SSAT-T’ and ‘Utility-T’ models for threshold parameters (R^2^=0.38) nor learning-rate parameters (R^2^=0.6).

**Table 2:** Estimated Models’ Parameters for Experiment 1 (Mean ± STD)

| Model | Beta | Learning Rate | Acceptability Threshold | Variance Learning Rate | Risk Aversion |
| --- | --- | --- | --- | --- | --- |
| Reward | 6.84 ± 3.67 | 0.61 ± 0.21 |  |  |  |
| Utility | 0.71 ± 0.38 | 0.55 ± 0.23 |  | 0.36 ± 0.15 | 0.91 ± 1.06 |
| SSAT | 5.65 ± 3.46 | 0.64 ± 0.21 | 0.52 ± 0.13 | 0.32 ± 0.21 |  |
| Reward-T | 8.48 ± 3.88 | 0.56 ± 0.19 | 0.33 ± 0.13 |  |  |
| Utility-T | 0.86 ± 0.45 | 0.49 ± 0.20 | 0.41 ± 0.17 | 0.39 ± 0.13 | 0.66 ± 0.63 |
| SSAT-T | 5.34 ± 2.37 | 0.58 ± 0.18 | 0.35 ± 0.13 | 0.31 ± 0.15 |  |

### Model predictions for confidence ratings

We hypothesize that confidence in choice reflects the subjective probability that the value of the chosen option exceeded the acceptability threshold (i.e., the total mass under the value distribution of the chosen option that is more than the acceptability threshold). To test this hypothesis, we compared the predictions of our models for confidence to the empirical confidence reports. We used the free parameters fitted to trial-by-trial choice data for each individual participant, and the values assigned to each option by the different models to draw predictions for the confidence reports for the corresponding individual. Following previous studies that examined confidence in value-based decisions [29,30], we defined confidence, for the ‘Reward’ and ‘Utility’ models, as proportional to the estimated decision variable: means of options’ rewards for ‘Reward’ models and the expected utilities of options for the ‘Utility’ models. We focused on trials 10-25 of each experimental condition and regressed the models’ predictions for these trials from the confidence reports made in these trials by each participant, in order to obtain the individual goodness of fit for each model (R^2^) (left column of Table T1, higher is better). We found that the model which gave the best predictions for trial-by-trial confidence reports was the ‘SSAT-T’ model, which formalized confidence reports as the probability of exceeding an acceptability threshold (paired t-test vs. Reward p = 0.04, vs. Reward-T p = 0.03, vs. Utility p = 0.005, vs. Utility-T p = 0.00002, vs. SSAT p = 0.16, Fig 3B).

To examine the pattern of confidence reports generated by each model, we calculated the average confidence for each model’s simulation when choosing the good option and when choosing the bad option in each condition. All models predicted lower confidence when choosing the bad, as compared to the good, option (Fig 4 and Fig S5 for the non-drifting models), similar to the observed confidence reports. Additionally, all models predicted similar confidence levels across conditions when the bad option was chosen. When choosing the good option, however, the SSAT-T model’s predictions were the most consistent with the reports made by participants. Only the SSAT-T model predicted higher confidence when variance of the good option was low (i.e. ‘vL-vL’,’vH-vL’). The other models did not predict a variance effect on confidence.

**Fig 4:**
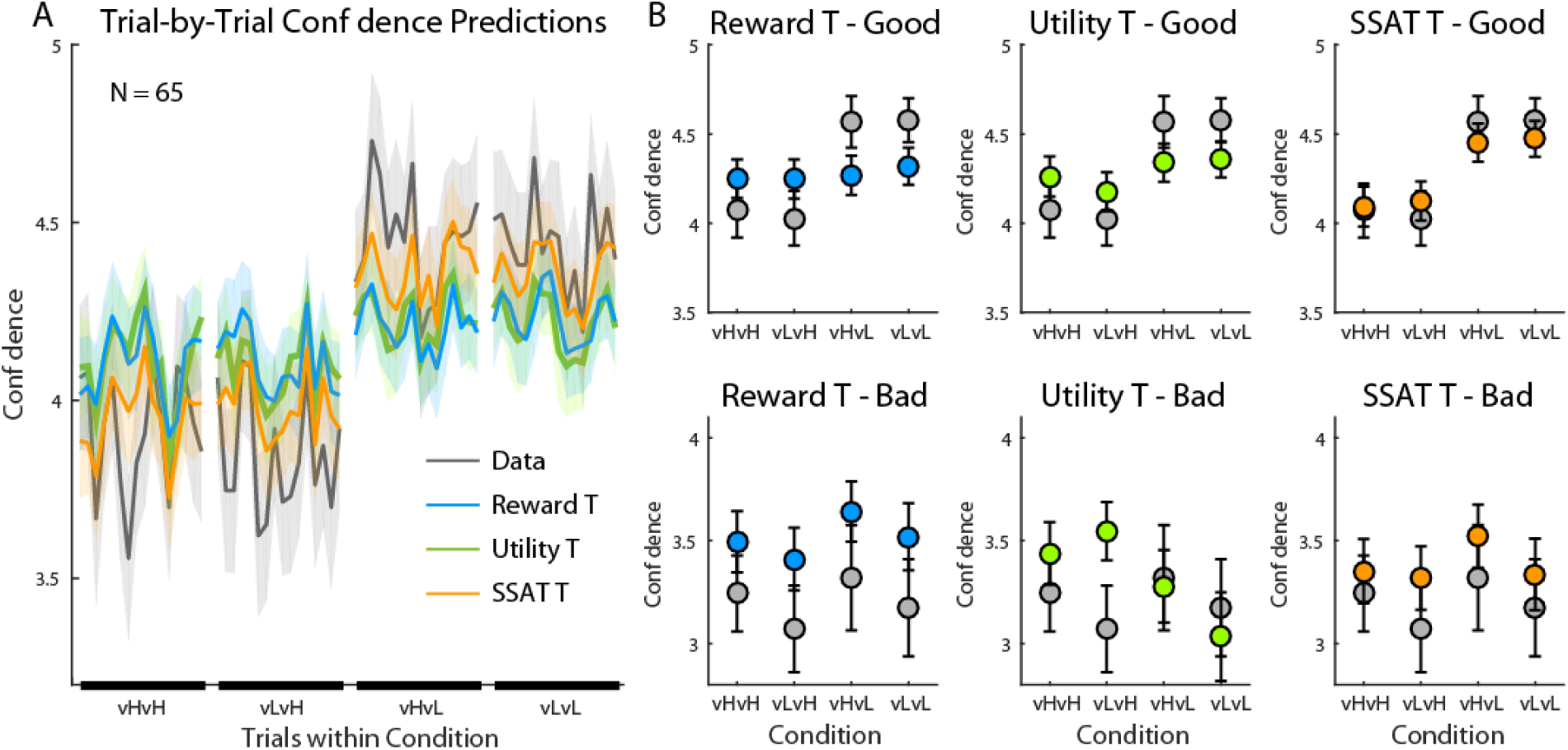
Model predictions for confidence reports in experiment 1. (A) Trial-by-Trial confidence reports averaged across participants (grey line) and model predictions during each experimental condition are displayed (shaded areas represent SEM). While the ‘Reward T’ and the ‘Utility T’ models gave similar confidence predictions across conditions, the ‘SSAT T’ model best corresponded with the data, as its confidence predictions increased when the variance of the good option was low. (B) Models’ predictions for confidence reports when choosing the good option (Top Row) and when choosing the bad option (bottom row). Predictions were averaged between trials 10-25 in each block. The average reports made by participants is displayed in grey. All models predicted higher confidence when choosing the good option than when choosing the bad option. ‘The SSAT T’ model gave the best predictions of confidence reports. Error bars represent SEM. The fit of the ‘Reward’, ‘Utility’ and ‘SSAT’ models are depicted in Fig S5.

Our model-comparison approach showed that the use of acceptability threshold parameter helped explaining participants’ choice behaviour, which was best captured by the ‘Reward-T’ model. Confidence, on the other hand, followed most closely the ‘SSAT-T’ model prediction of reporting the probability of exceeding the acceptability threshold. A counterintuitive prediction of the model borne out by the behavioural data was the difference between the two conditions involving unequal variances (i.e. vL-vH and vH-vL conditions). Stochastic satisficing predicted – and the data confirmed – a difference in confidence (c.f. Fig 4, compare vL-vH and vH-vL) despite identical expected values for the chosen (good) option in these two conditions. In Experiment 2, we focused on the choice between options with unequal variances to further tease apart the cognitive substrates of stochastic satisficing.

### Experiment 2

To conduct a more rigorous test of the parsimony and plausibility of the stochastic satisficing heuristic, in Experiment 2, we designed a new payoff structure for the two-arm bandit, focusing on options with unequal variances in all conditions (Fig 5A). We kept the mean and variance of the bad option constant across conditions (mean=35 and variance=10^2^=100) while varying the mean and variance of the better option in a 2 × 2 design. Mean reward of the good option could be low (mL=57; still better than the bad option) or high (mH=72), and its variance could be independently low (vL=5^2^=25) or high (vH=20^2^=400). Thus, we constructed four experimental conditions all involving options with unequal variances and large or small differences in their expected values. We followed the optimality analysis described above with the reward distribution of Experiment 2, and found that our models predicted a distinctive and different pattern of confidence in each condition of the new design (Fig S7).

**Fig 5:**
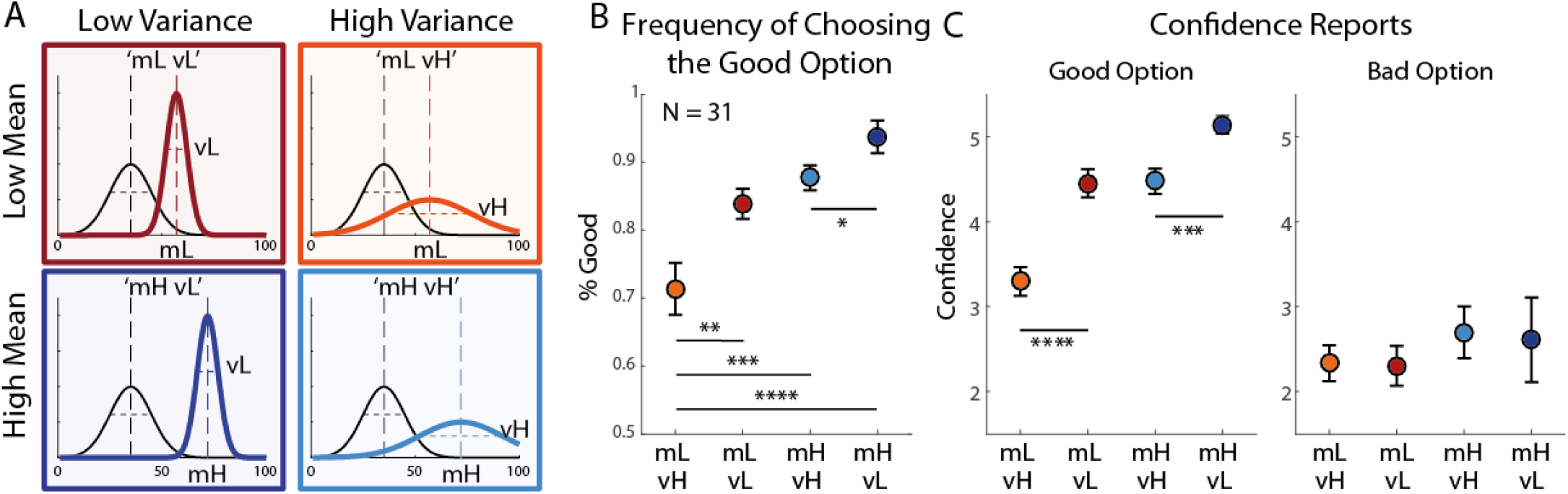
Experiment 2 design and behavioural results. (A) In experiment 2 the rewards’ mean and variance of the bad option (black lines) were kept constant across experimental conditions, while the mean and variance of the good option varied. Mean values could be high (mH) or low (mL), and variances could be independently high (vH) or low (vL), resulting in four experimental conditions. (B) Experimental results (33 subjects). Both choices and confidence reports were averaged between trials 10 to 25 of each experimental block. Frequency\ of choosing the good option gradually increased as the mean expected reward increased, and as the variance decreased. (C) When choosing the good option (middle panel), confidence ratings did not differ between the ‘mL-vL’ and ‘mH-vH’ condition. When choosing the bad option (right panel) confidence reports were not significantly different between conditions. Error bars represent SEM. (* p<0.05, *** p<0.0005).

We examined choices and confidence reports in experiment 2 in a new group of participants (N=31). The probability of choosing the good option increased with the mean reward of the good option (mixed effects ANOVA, F(1,89)=21.25, p=0.0001) and decreased with its variance (F(1,89)=15.03, p=0.0005) (Fig 5B). Confidence when choosing the bad option did not change significantly across conditions (Fig 5C). When choosing the good option, confidence was significantly affected by variance (F(1,89)=35, p<0.00001) and mean (F(1,89)=88, p<0.00001) of the good option’s rewards. However, while confidence in the ‘mH-vL’ was significantly higher than all other conditions (‘mH-vL’ vs ‘mH-vH’: t(30)=4, p=0.0003), and confidence in the ‘mL-vH’ was lower than all other conditions (‘mL-vH’ vs. ‘mL-vL’: t(30)=4.9, p=0.00002), the critical comparison of ‘mL-vL’ and ‘mH-vH’ did not show a difference (‘mL-vL’ vs. ‘mH-vH’: t(30)=0.4, p=0.68). Finally, we examined the relations between confidence report and reaction time and found that the participants’ correlation coefficients were significantly lower than zero (t-test, p=0.0014, Fig 6). However, just like in Experiment 1 the overall link across participants between confidence and reaction time was not very strong, with average correlation coefficient of R = −0.067, below the critical value of R(160) = 0.16 for significance of 0.05. We examined the relationship between confidence reports and choice stability, and found that the participants’ correlation coefficients were significantly higher than 0 (t-test, p< 10-^13^, Fig 6), with average correlation coefficient of R = 0.44.

**Fig 6:**
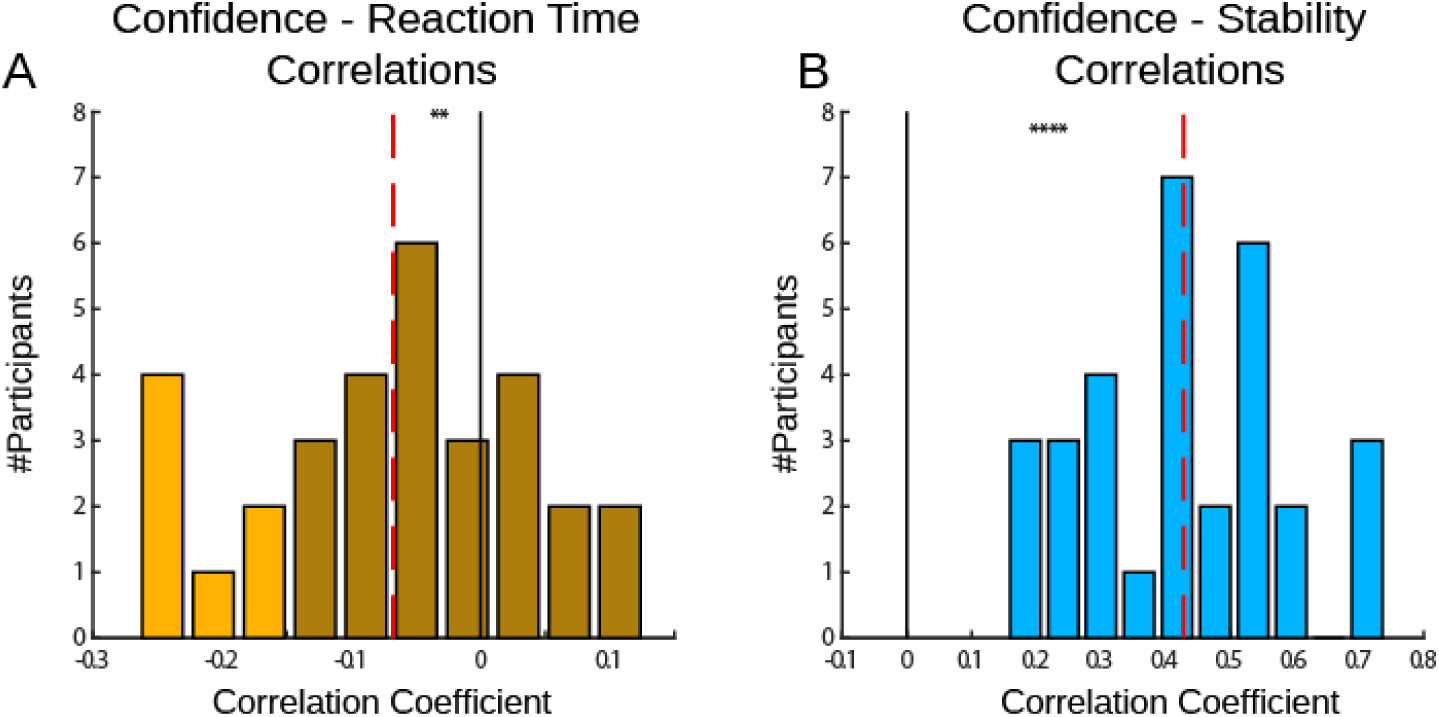
Correlations between confidence, reaction time, and stability in Experiment 2. (A) We correlated each participant’s reaction times with confidence ratings. We found that the participants’ correlation coefficients tended to be below 0, as fast responses were associated with higher confidence. However, these were not as strongly linked across the population, with average correlation of R= −0.067 (dashed line). (B) We examined the relations between confidence and choice stability. We found that the participants’ correlation coefficients were highly significant in the individual level and in the group level (average R = 0.44, dashed line). Dashed red lines indicate the average correlation coefficient. Dark colours indicate below significance correlation (Critical value of R(160) = 0.16, p = 0.05). ** p < 0.005, **** p < 0.00005.

### Model-fitting and Predictions

We fitted all models to the choices made by participants in Experiment 2. Like in Experiment 1, adding the acceptability threshold parameter helped explaining participants’ choice behaviour in all models (Table 3). We found again that the best description of the data was given by the ‘Reward-T’ model, however not as strongly as in experiment 1 (paired t-test T vs. Reward p = 0.25, vs. Utility p = 0.07, vs. Utility-T p = 0.05, vs. SSAT p = 0.28, vs. SSAT-T p = 0.06, Fig 7A). When examining the relationship between individual parameters fitted by the models, we found again a high correspondence between the parameters estimated for Acceptability Threshold and Learning Rates between the ‘Reward-T’ and ‘SSAT-T’ model (Table 4), captured by high correlation between the individual threshold parameters (R^2^=0.9) and learning rate parameters (R^2^=0.82) (Fig S8). Such similarity was not found between the ‘SSAT-T’ and ‘Utility-T’ models for neither threshold parameters (R^2^=0.15) nor learning rate parameters (R^2^=0.34).

**Table 3:** Models performance in Experiment 2.

| Model | Sum WAIC | Mean ± STD WAIC | Mean ± STD Confidence $R^2$ |
| --- | --- | --- | --- |
| Reward | 3,971 | 128.09 ± 43.39 | 0.18 ± 0.15 |
| Utility | 3,803 | 122.69 ± 35.86 | 0.34 ± 0.19 |
| SSAT | 3,957 | 127.66 ± 44.48 | 0.21 ± 0.18 |
| Reward-T | **3,629** | 117.09 ± 33.31 | 0.35 ± 0.15 |
| Utility-T | 3,770 | 121.62 ± 35.46 | 0.31 ± 0.13 |
| SSAT-T | 3,703 | 119.48 ± 33.62 | **0.41 ± 0.18** |

**Table 4:** Estimated Models’ Parameters for Experiment 2 (Mean ± STD)

| Model | Beta | Learning<br>Rate | Acceptability<br>Threshold | Variance<br>Learning<br>Rate | Risk Aversion |
| --- | --- | --- | --- | --- | --- |
| Reward | 9.20 ± 3.73 | 0.56 ± 0.20 |  |  |  |
| Utility | 1.05 ± 0.52 | 0.54 ± 0.19 |  | 0.31 ± 0.13 | 1.00 ± 0.95 |
| SSAT | 5.22 ± 2.9 | 0.61 ± 0.21 | 0.51 ± 0.79 | 0.42 ± 0.15 |  |
| Reward-T | 12.17 ± 3.72 | 0.54 ± 0.17 | 0.38 ± 0.11 |  |  |
| Utility-T | 1.76 ± 1.4 | 0.43 ± 0.17 | 0.51 ± 0.15 | 0.36 ± 0.15 | 0.69 ± 0.59 |
| SSAT-T | 6.54 ± 2.25 | 0.51 ± 0.17 | 0.41 ± 0.09 | 0.33 ± 0.21 |  |

**Fig 7:**
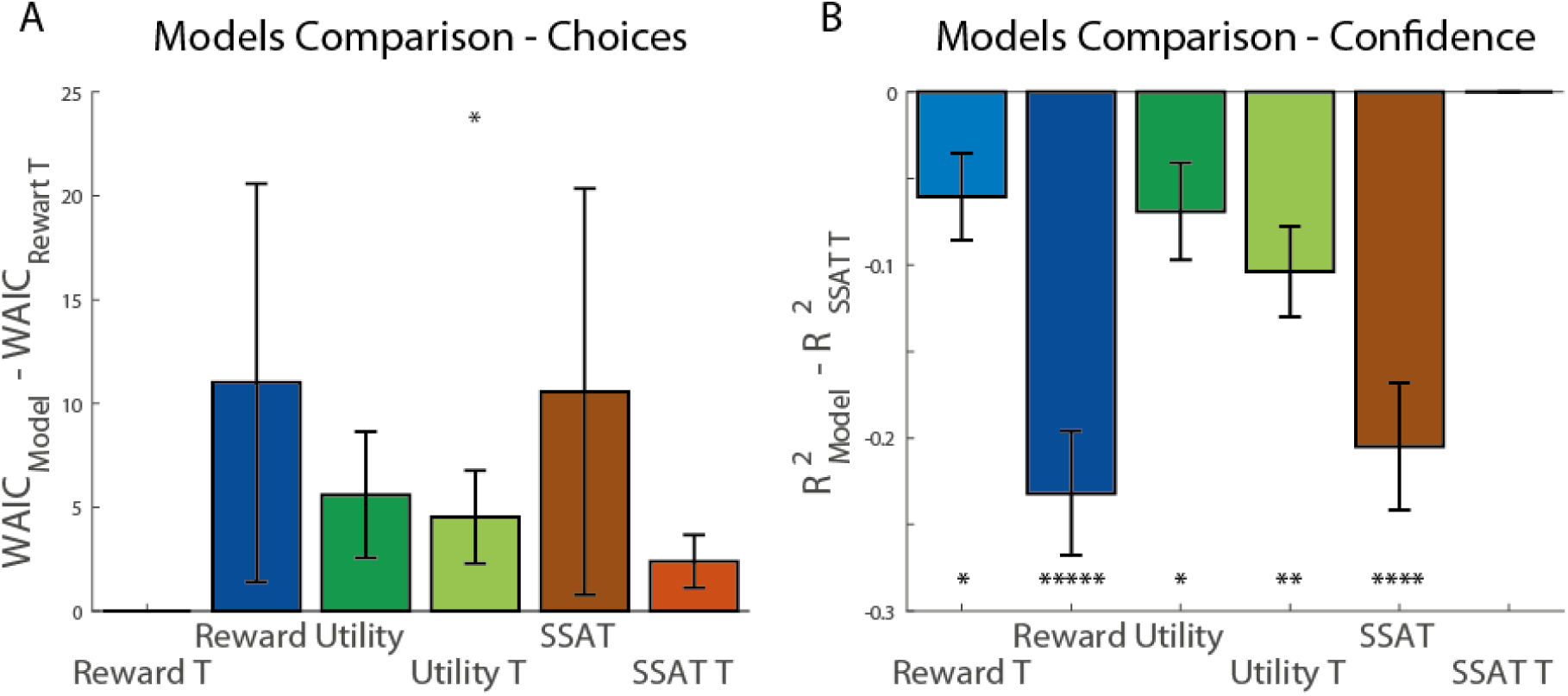
Models comparison in experiment 2. (A) We compared the ‘Reward T’ (lowest WAIC) model WAIC score to the other models by examining the paired differences in WAIC scores across models and participants. The graph presents the differences of each model WAIC from that of ‘Reward T’ model. The ‘Reward T’ model performance was not significantly better than most of the other models in explaining participants’ choices. (B) We compared ‘SSAT T’ model (highest R^2^) to all other models by examining the paired differences in R^2−^scores across models and participants. The ‘SSAT T’ model gave a significantly better prediction of confidence reports than all other models. Error bars represent SEM. * p<0.05, ** p<0.005, *** p<0.0005, **** p<0.00005.

Experiment 2 was explicitly designed to test the models’ predictions of choice confidence. We examined whether the models estimated trial-by-trial decision variables (means of rewards for ‘Reward’ models, expected utility for the ‘Utility’ models, and probability of exceeding acceptability threshold for the ‘SSAT’ models). Just like in Experiment 1, we focused on trials 10-25 of each experimental condition, and regressed the models’ predictions from these trials from the confidence reports made in these trials for each participant, to obtain the individual goodness of fit for each model (R2) (right column of Table 3, higher is better). We found that the model which gave the best predictions of trial-by-trial confidence reports in Experiment 2 was the ‘SSAT-T’ model (paired t-test vs. Reward p = 0.0000003, vs. Reward-T p = 0.02, vs. Utility p = 0.02, vs. Utility-T p = 0.0004, vs. SSAT p = 0.00004, Fig 7B).

To demonstrate the pattern of confidence reports generated by each model, we calculated the average confidence for each model’s simulation when choosing the good and the bad options in each condition. The most striking qualitative difference between the models was in their predictions of confidence reports when choosing the good option. All models predicted the lowest confidence for choosing the good option with low mean and high variance (‘mL-vH’) (Fig 8 and Fig S9). Highest confidence was predicted when choosing the good option with high mean and low variance (‘mH-vL’) by all models. However, ‘SSAT-T’ model was the only one following the pattern observed in the participants’ reported confidence, predicting similar confidence ratings for the low mean, low variance (‘mL-vL’) and the high mean, high variance (‘mH-vH’) conditions, as the probability of exceeding the satisficing threshold was the same for these two conditions. Critically, because these two conditions had different expected rewards, the ‘Reward-T’ model predicted different confidence levels for them. Even though ‘Utility-T’ model penalized options’ values according to their variance, it failed to recover the behavioural pattern and predicted lower confidence reports in the ‘mL-vL’, compared to the ‘mH-vH’, condition.

**Fig 8:**
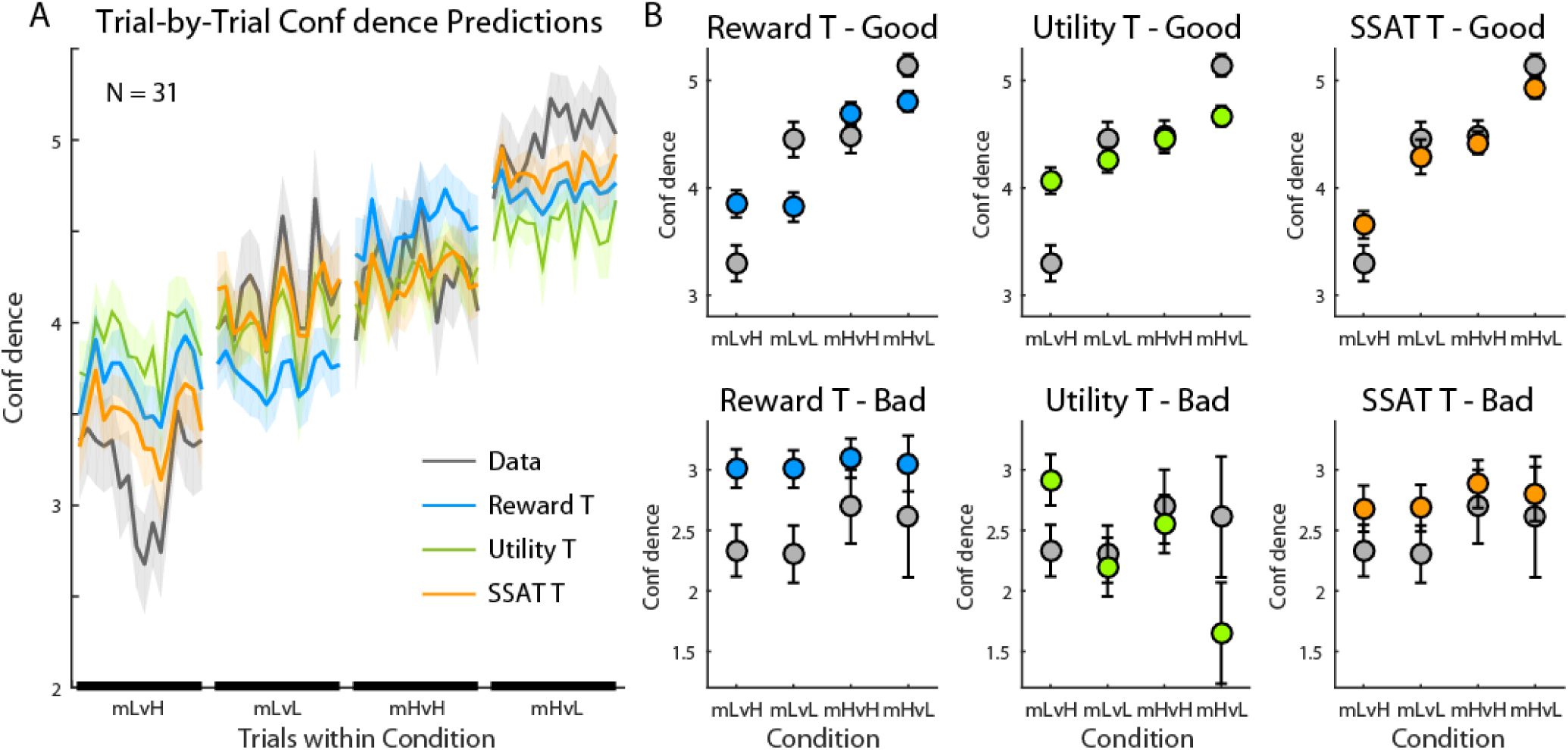
Model predictions for confidence reports in experiment 2. (A) Trial-by-Trial confidence reports (grey line) and model predictions during each experimental condition are displayed, averaged across participants (shaded areas represent SEM). The ‘SSAT T’ model best corresponded with the data, as its confidence predictions were dependent on both the mean and the variance of the reward distributions. (B) Models’ predictions for confidence reports when choosing the good option (Top Row) and when choosing the bad option (bottom row). Predictions were averaged between trials 10-25 in each block. The average reports made by participants is displayed in grey. SSAT-T model gave the best prediction of confidence reports. Error bars represent SEM. The fit of the ‘Reward’, ‘Utility’ and ‘SSAT’ models are depicted in Fig S9.

## Discussion

We set out to examine decision-making and confidence reports in uncertain value-based choices. In a two-armed bandit task played by human participants, the probability of choosing the good option increased as the variance of either options’ outcomes decreased. However, confidence ratings were associated with variance only when choosing the good (higher mean) option, as items with low variance outcomes were chosen with higher decision confidence. Confidence ratings associated with choosing the bad (i.e. lower mean) option were always low and were independent of the variances of the options’ outcomes. We examined how bounded rationality heuristics may account for this pattern of behaviour, first by introducing an acceptability threshold representing an expectation about the outcomes’ values, and by proposing a stochastic satisficing model in which decisions are made by comparing the options’ probability of exceeding this acceptability threshold [31]. We found that choice behaviour could be accounted for by adding a threshold parameter to a simple TD learning mechanism which tracks the expected reward of each option [21,25]. Confidence reports, however, were best captured by the stochastic satisficing model, as confidence reports scaled with the chosen option’s satisficing probability. To directly test a critical prediction of this model, a second experiment involving options with unequal variances and means was simulated first and then empirically performed. As predicted by the ‘SSAT-T’ model, participants’ confidence reports matched the options’ probability of exceeding a threshold, and not the options’ expected outcome.

In our experiments, models aimed at maximizing expected utility [4,22], modelling the impact of risk aversion on options’ values, were not successful at explaining participants choices or confidence reports. Maximizing the expected exponential utility function boiled down to penalizing outcomes according to their variance (i.e. Mean-Variance paradigm, see Methods and [23,24]). An important feature of this instantiation of risk aversion is that the effect of variance is always in the same direction, reducing the value or utility of both good and bad options. This means that when the variance of the bad option increases, the likelihood of choosing the good option should increase. This was not the case in our experimental results. Our stochastic satisficing model provides a mechanism by which variance effect is not symmetrical for good and bad option – when the bad option’s variance increases, its value (i.e. the probability of surpassing an acceptability threshold) increases.

Our results suggest a divergence between choice and confidence reports. Choices were best explained by the ‘Reward-T’ model, which does not track outcome’s variance, while confidence reports were best explained by the ‘SSAT-T’ model and were affected by the outcomes’ variance. Our optimality analysis also demonstrated this separation between the performance of a ‘greedy’ decision-maker, insensitive to the size of the value-difference between options, and a ‘noisy’ decision-maker whose likelihood of choosing the high-value option, as well as its decision confidence, scale with the amount of evidence favouring that option. Such separation of actions and evaluation of actions is in line with the second-order framework for self-evaluation of decision performance [18]. In the second-order model suggested by Fleming and Daw, action and confidence stem from parallel processes. Sensory input is assumed to be sampled independently by the action and evaluation processes. In this framework, confidence is first affected by its independent sample, and then by the action chosen by the action process. Our results are in line with such parallel processing. Actions were accounted for by a parallel and correlated process to the confidence generating process, and confidence was conditional on the action selected by the client.

The second-order model [18] and other recent studies [14,17,19,32] have formulized confidence as the probability of having made a correct choice over tracked outcome or evidence distribution. This approach builds on the line of research about the representation of evidence distribution, and suggests that confidence summarizes this probabilistic representation, estimating the probability of being correct. Probability of being correct is more readily defined in perceptual detection tasks where option outcomes are not independent (e.g. the target can be in only one of two locations but not both) and there is an objective criterion for correctness. Our stochastic satisficing model expands these observations from perceptual decisions to scenarios where outcomes are stochastic. In such scenarios, our theory-based analysis of data suggests, participants use an arbitrary criterion, the acceptability threshold, to evaluate the probability of an outcome to exceed the threshold, analogous to the evaluation of correctness probability in detection tasks. Confidence would then reflect the probability that the chosen option exceeded the “good enough” acceptability threshold. As the likelihood of exceeding the acceptability threshold increases – either by reducing the outcome variance (Experiment 1) or increasing the outcome mean (Experiment 2) – so does decision confidence.

Another important divergence of our design from perceptual decision tasks is the relatively weak link between reaction time and confidence reports. In perceptual decision making, the entire process of evidence accumulation is encapsulated in one trial and a drift diffusion model can therefore capture this process and predict response time and confidence at the same time [16,29]. In our learning task, evidence about each option’s reward distribution was accumulated across trials – on each trial, the participant sampled one option and learned from its reward. In this case, the link between reaction time and confidence reports may not be as strong as in the perceptual tasks. Our ‘choice stability’ measure was found to be highly correlated with confidence reports. This measure can be interpreted as an indication of how many favourable examples the participant accumulated before making the confidence report.

This process is similar to the evidence accumulation process modelled by the drift diffusion model, but in our case the accumulation is across trials and not within a trial. This discrimination is important and may shed light on evidence accumulation process in the brain. Our design provides an opportunity for future research on the neural mechanism of metacognition, as it integrates previous knowledge about representation of variance [33,34] in the brain with the literature on neural mechanism of metacognition [15,35], and allows a better dissociation of decision and confidence in the brain.

In the 1950s Simon introduced the concept of satisficing, by which decision makers settle for an option that satisfies some threshold or criterion instead of finding the optimal solution. The idea is illustrated in the contrast between ‘looking for the sharpest needle in the haystack’ (optimizing) and ‘looking for a needle sharp enough to sew with’ (satisficing) (p. 244) [12,36]. This notion of acceptability threshold has been extended to other ambiguous situations [37], for example for setting a limit (i.e. threshold) to the time and resources an organization invests in learning a new capability [12], where suboptimal solution may be balanced with preventing unnecessary cost. We suggest that stochastic satisficing serves a similar objective by extending the basic idea of satisficing into stochastic contexts with continuous payoff domains [13]. We found stochastic satisficing to be particularly useful at explaining decisions’ confidence, i.e. evaluation of decisions. Bounded rationality was originally developed to explain administrative decision making [11], in which decisions are often evaluated explicitly, and the decision maker is held accountable for the outcome [38]. Stochastic satisficing may therefore serve psychological and social purposes associated with the evaluation, communication and justification of decision-making [39,40]. As it strives to avoid catastrophe, i.e. receiving a reward below acceptability threshold, stochastic satisficing may be useful to minimize regret, similarly to status quo bias [41,42]. Choosing the option less likely to provide unacceptable payoffs can serve as a safe argument for justifying decisions to oneself or others [38], in the spirit of the saying “nobody ever got fired for buying IBM”.

## Methods

### Participants

We recruited participants through Amazon M-Turk online platform [43]. All participants provided an informed consent. Experiments were approved by the local ethics committee. Participants earned a fixed monetary compensation, but also a performance-based bonus if they collected more than 10,000 points. 88 participants were recruited for the first experiment, in order to obtain power of 0.8 with expected effect size of 0.4 for variance effect on confidence. The actual effect size obtained in Experiment 1 was 0.6, and we therefore recruited 33 subjects for the second experiment. 25 participants were excluded from analysis as their performance was at chance level (16 participants) or for using only one level for confidence reports (9 participants). Data from 96 participants (62 males aged 32±9 (mean±std), and 34 females aged 32±8) were analysed.

### Experimental Procedure and Design

On each trial participants chose between two doors, each leading to a reward between 1 and 100 points (Fig 1A). Each door had a fixed colour-pattern along the task, but the positions (left vs. right) were chosen randomly. Subjects made choices by using a 12-level confidence scale: 1-6 towards one option and 1-6 towards the other, with 6 indicating ‘most certain’ and 1 indicating ‘most uncertain’. Following choice, subjects observed the reward of the chosen door drawn from a normal distribution 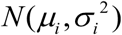, where *i* was *a* or *b*, indicating one or the other door. A working demo of the task can be found here: http://urihertz.net/BanditConfDemo/.

Experiment 1 consisted of 240 trials and included six stable blocks where the mean and variance of each option’s reward remained constant. Each block lasted at least 25 trials. The transition from one block to another occurred along 10 trials during which the mean and variance associated with each door changed gradually in a linear fashion, from their current to the new levels corresponding to the upcoming block. Embedded within these six blocks, four blocks followed a 2 × 2 design where the mean rewards of the two options were 65 (for the good option) and 35 (for the bad option), and their variances could be independently high (H=25^2^=625) or low (L=10^2^=100) (Fig 1B). This design included four conditions: ‘vL-vL’,’vH-vL’,’vL-vH’ and ‘vH-vH’, where the first and the second letters indicated the magnitude of the variance of the good and the bad options, respectively.

Experiment 2 consisted of 160 trials and was similarly composed of blocks of fixed reward probability distributions. In all four blocks, the reward of one option always followed a Gaussian distribution with a mean of 35 and a variance of 100 (10^2^). The mean of the other option could take either high (mH=72) or low (mL=57), and its variance could be either high (vH =20^2^=400) or low (vL=5^2^=25). This produced a 2 × 2 design, with the four conditions denoted by ‘mL-vL’, ‘mL-vH’, ‘mH-vH’, ‘mL-vL’(Fig 5A).

## Models

Six different models were fitted to the participants’ choices. These included models that track only mean of the rewards from each option, and models that track both mean and variance.

The ‘Reward’ model assumes that expected reward of the outcomes govern choices, and it tracks the means of the rewards using a temporal difference algorithm [21,25].

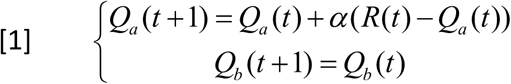

Where a and b indicate the chosen and the un-chosen options, respectively. α is the learning rate. A softmax action-selection rule was used:

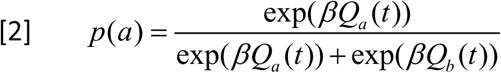

Where β is the rate of exploration. Therefore, the ‘Reward’ model has 2 free parameters: *{α, β}.*

The ‘Utility’ model tracks both mean and variance of rewards from the two options. Tracking the mean of rewards is done in a similar manner to the ‘Reward’ model (Eq. [1]). Tracking of variance is done using a similar temporal difference rule:

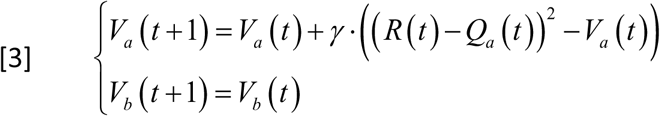

Where γ is the variance learning rate. This model assumes an increasing and concave exponential utility function [5,24]by which the utility of a reward decreases as the reward increases:

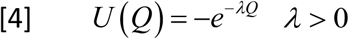

*λ* denotes the risk sensitivity of the participant, the larger *λ* is, the more risk averse the participant is. When rewards are governed by a Gaussian distribution is it possible to evaluate the expected utility of an option analytically [22,24].

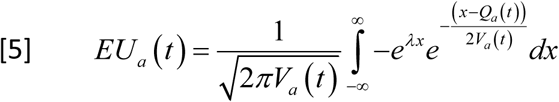

Using the tracked variance and mean of the rewards, the value to maximize (for option a, for example) is:

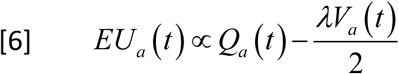

This is a formulation of the variance-mean balance, in which choices’ expected utility depends on the expected reward, and penalized by the variance of rewards [23,24]. A softmax rule (Eq. [2]) was used for action selection with the expected utilities as the values associated with each option. The ‘Utility’ model has 4 free parameters: *{α, γ, λ, β}.*

The Stochastic satisficing (SSAT) model employs a threshold heuristic [13]. It tracks the means (Eq. [1]) and variances (Eq. [3]) associated with the two options. The probability of payoff being higher than the acceptability threshold, *T*, is calculated using a cumulative Gaussian distribution equation:

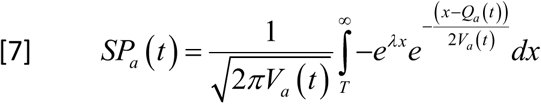

where SP_a_ indicated the probability of action *a* being satisficing. A softmax (Eq. [2]) rule is used to calculate choice probabilities according to the options’ satisficing probabilities (SPa and SP_b_).

In addition to these three models, we tested a version of all three models in which the unchosen option (in the example below option b was not chosen) drifts towards an acceptability threshold T:

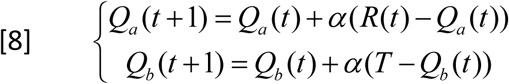

This rule was use in the ‘Reward-T’ and ‘Utility-T’ models, adding to them an additional free parameter T. This rule was also used in the ‘SSAT-T’ model, using the threshold parameter T which was already used in the ‘SSAT’ model.

## Optimization Analyses

We carried two analyses to examine the optimal performance our models are capable of in our experimental design, in terms of amount of reward accrued. In the first analyses we used parameter estimation (Nelder-Mead algorithm implemented by Matlab’s fminsearch function) to identify a set of parameters that maximizes each model’s accumulated reward. We then simulated the model choices using the identified parameters, and tracked how much reward was collected by each model over 100 repetitions.

In the second optimality analyses we set the reward distribution parameters, and examined the differences in values assigned to each option in each experimental design by the different models. We changed the value of the acceptability threshold, and examined how it affected the model’s estimations. This analysis allows a detection of the pattern of confidence and choice probabilities across conditions during steady state – after the reward distributions were learned.

### Model fitting and model comparison

For each model, we used Hamiltonian Monte Carlo sampling implemented in the STAN software package [44] to fit the free parameters of each model to the choice data, in a subject-by-subject fashion, in order to maximize likelihood [45]. The Markov Chain Monte Carlo (MCMC) process used for optimization produces a likelihood distribution over the parameter space of the model, for each subject. For model comparisons, we calculated Watanabe Akaike Information Criterion (WAIC) that uses these likelihood distributions and penalizes for the number of free parameters [28]. We then simulated each model, using the estimated posterior distribution over the individual parameters of that model, in order to produce the model’s value estimations associated with each option during the experiment. We used these estimated values to predict the model’s confidence ratings for the choices made by participants.

## Data Availability

The behavioural data and scripts supporting the findings of this study are available from Figshare (https://figshare.com/s/a6088b11227d9e61680c).

## Acknowledgements

UH and BB are supported by the European Research Council (NeuroCoDec 309865). UH is also supported by the John Templeton Foundation. MK is supported by the Gatsby Charitable Foundation.

## Supplementary Information

**Fig S1.**
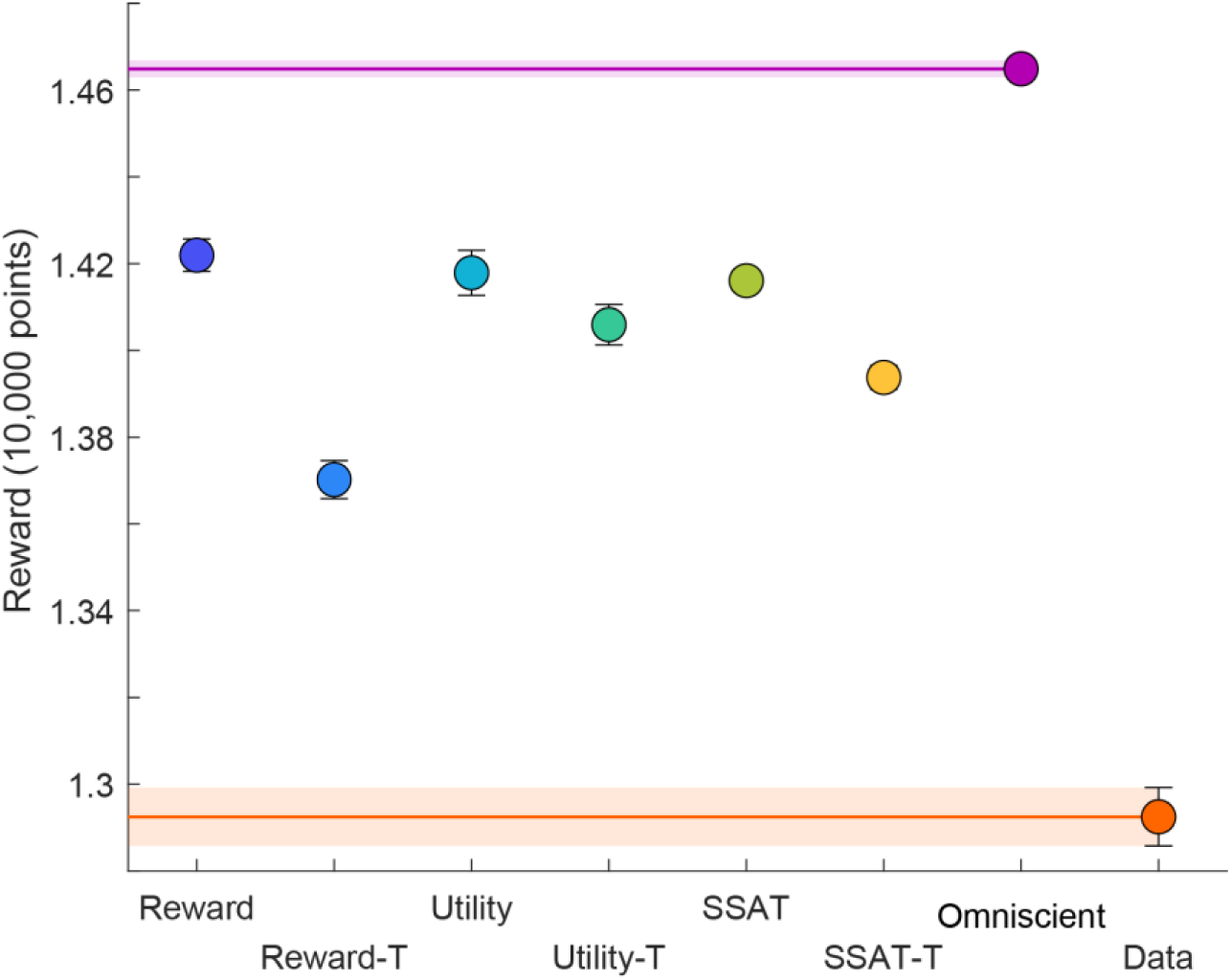
Trial-by-Trial Optimality Analysis Experiment 1. We identified parameters that maximized the amount of reward accumulated by each of our models with the reward distribution from experiment 1, and examined the amount of reward collected by the models using these parameters over 100 repetitions. We also examined the rewards accrued by a model with full knowledge of the reward distribution (‘Omniscient’), and the actual amount of reward accrued by our participants. We found that all three models performed similarly, and accrued similar amount of reward. When the drifting mechanism was added (drift of the unchosen option towards the acceptability threshold) performance of all models decreased. All models did not accrue as much reward as the ‘Omniscient’ model, as all of them had to learn and adapt to a dynamic and changing environment. In addition, all models performed much better than our participants, indicating that participants’ behaviour was noisy, falling short of the optimal strategy.

**Fig S2.**
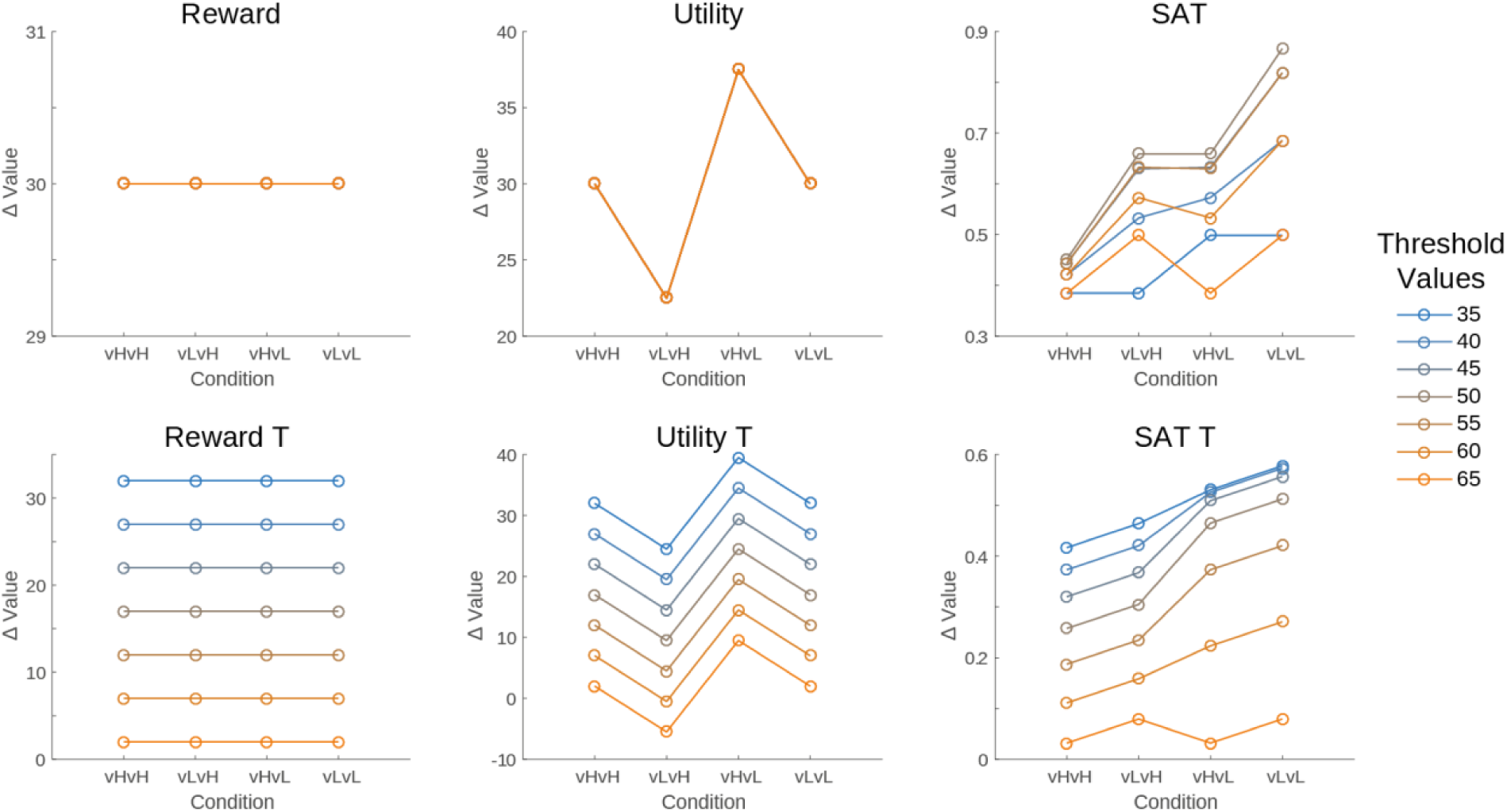
Value-Differences Optimality Analysis in Experiment 1. We examined the differences in values assigned to the two options by each model in the four conditions of our experimental design. We used the reward distributions mean and variances in each condition, and varied the acceptability threshold (blue to yellow lines). We found that all models assigned higher values to the high mean reward option than to the low mean reward option in almost all the cases and conditions. A greedy decision maker would therefore be able to accumulate similar amount of rewards using each model. However, different models assigned different value differences in each condition. This means that a noisy decision maker (modeled using softmax) may be more likely to choose the low mean reward option in some conditions, according to the models’ predictions.

**Fig S3.**
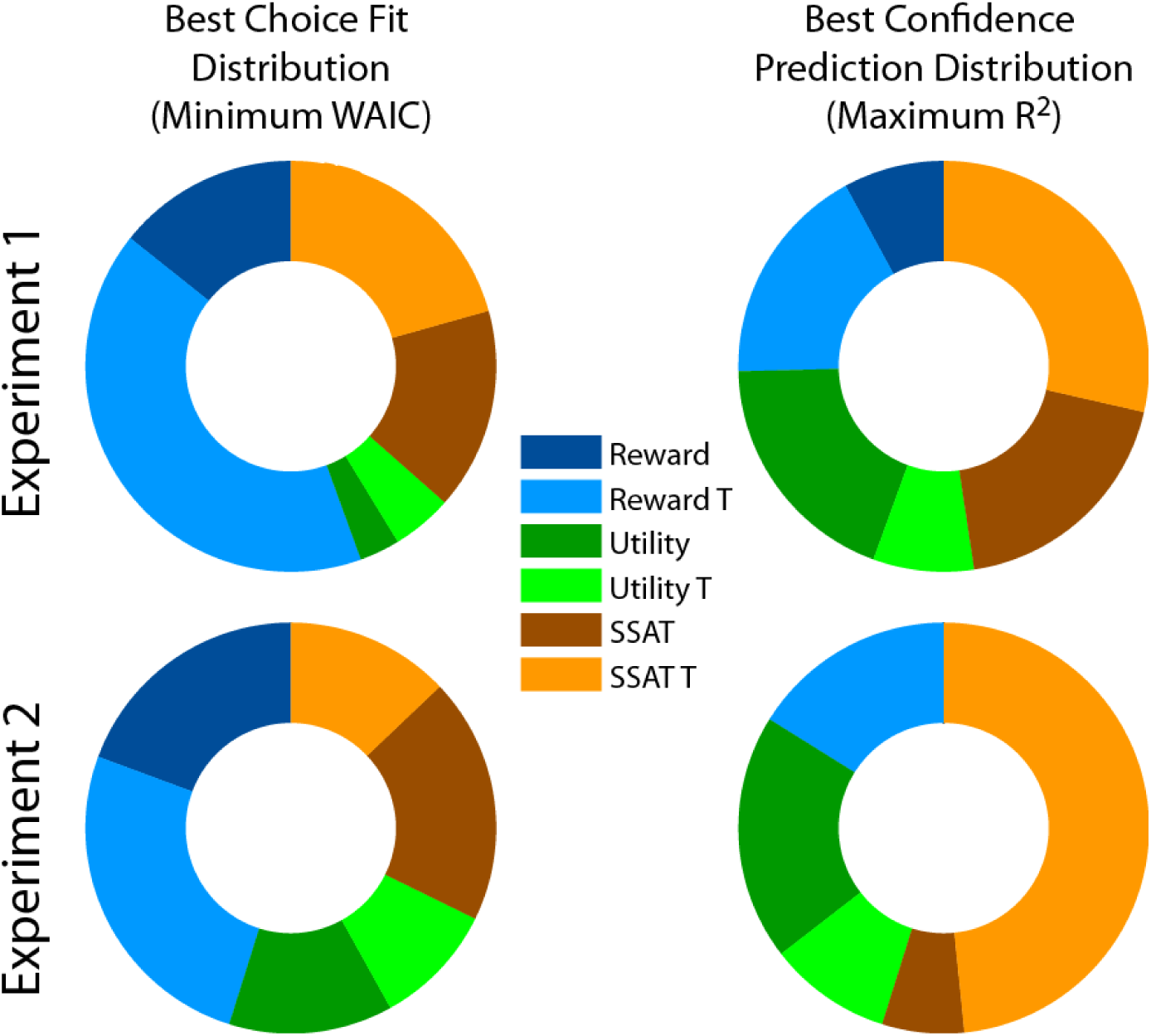
Distribution of Best Model Fits Across Participants. We examined how many of the participants’ choices were best explained by each of our six models in both experiments (left panels), and how many participants’ confidence reports were best predicted by the models (right panels). We found that in Experiment 1 most of the participants’ choices were best explained by models that did not track reward variance, in line with the model comparisons we performed. In Experiment 2 choice responses were split between models that tracked variance and models that did not track variance. Best confidence ratings predictions were also distributed across participants. We found that in Experiment 1 most participants’ confidence reports were affected by variance, with half of the participants’ confidence reports best predicted by the SSAT or SSAT-T models. In Experiment 2 the picture was even more robust, with even greater share of the participants’ reports being affected by outcome variance. The distributions of confidence and choices were found to be different (Two-sample Kolmogorov-Smirnov test, Experiment 1: p = 0.0049, Experiment 2: p = 0.03).

**Fig S4.**
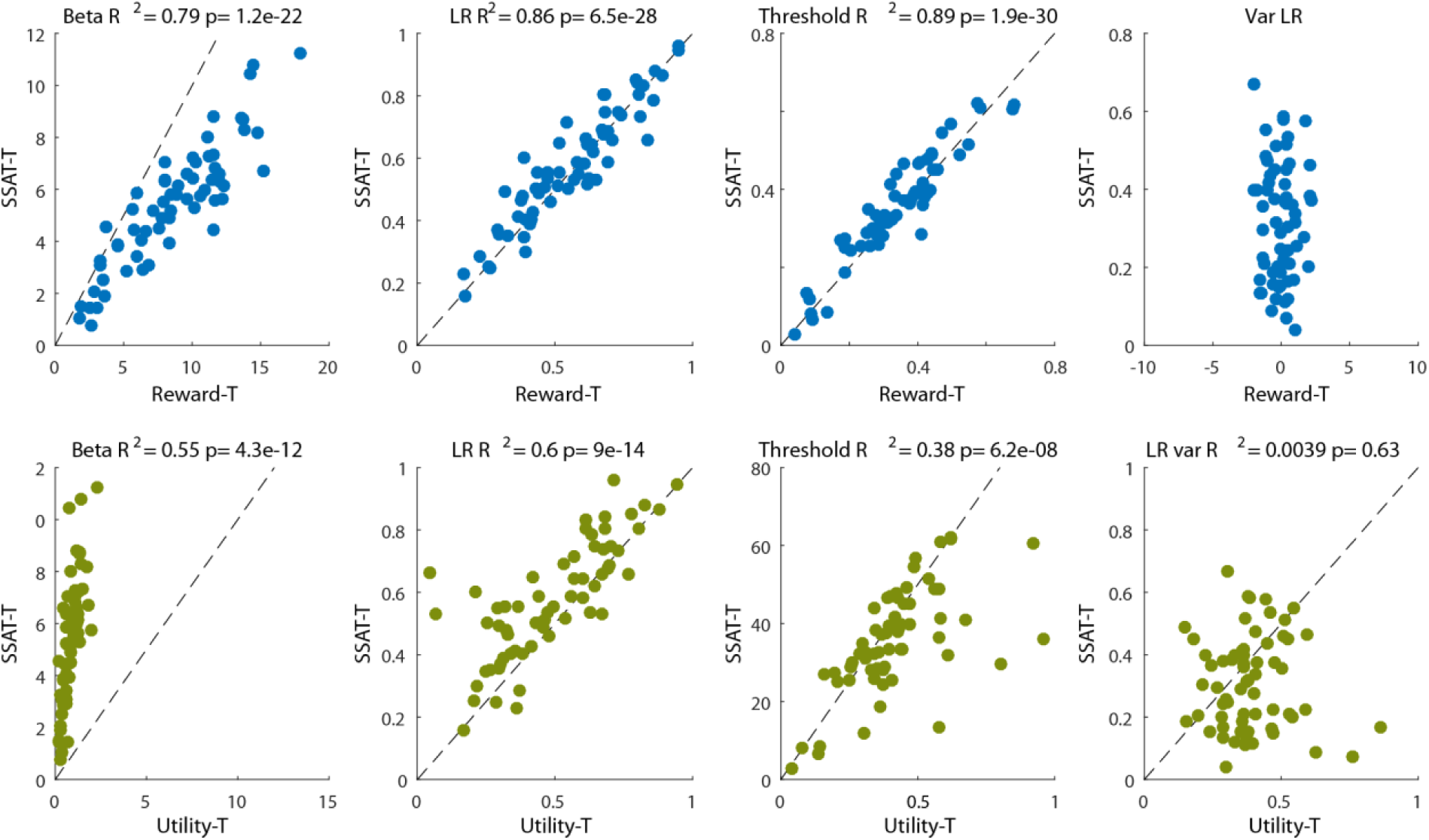
Relations between estimated parameters in different models in experiment 1. We compared the individual parameters estimated for each of our drift (and best performing) models. The Reward-T and SSAT-T models’ parameters for learning rates and threshold were almost identical for all our participants. Reward model gave the best fit to decisions, while the SSAT-T model gave the best fit to confidence reports. This indicates that these models may use a shared mechanism for decisions, but the SSAT-T model uses the reward variance information to generate confidence reports. Parameters estimations were not as similar for the SSAT-T and the Utility-T.

**Fig S5.**
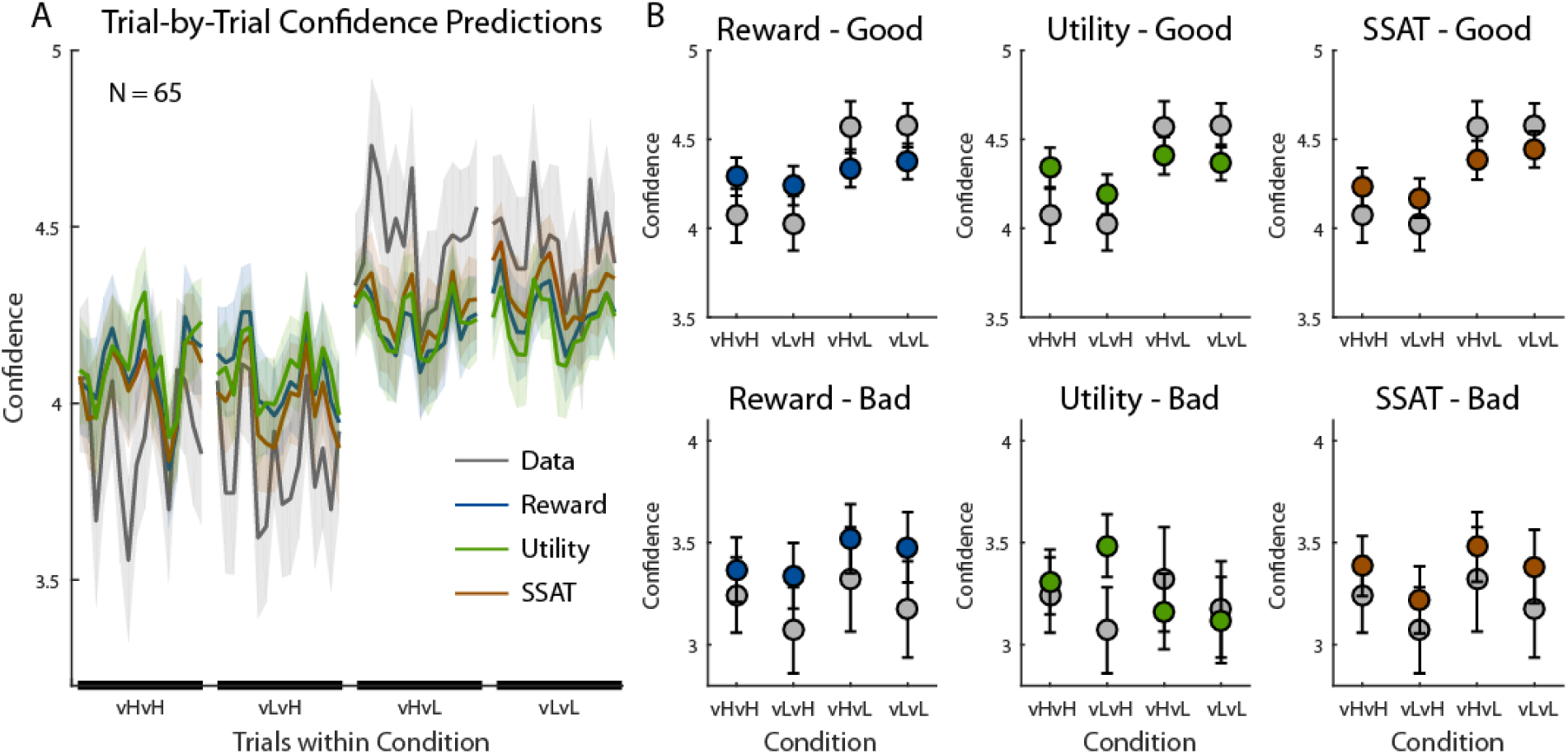
Model predictions for confidence reports in experiment 1. (A) Trial-by-Trial confidence reports (grey line) and model predictions during each experimental condition are displayed, averaged across participants (shaded areas represent SEM). (B) Models’ predictions for confidence reports when choosing the good option (Top Row) and when choosing the bad option (bottom row). Predictions were averaged between trials 10-25 in each block. The average reports made by participants is displayed in grey. All models predicted higher confidence when choosing the good option than when choosing the bad option. Error bars represent SEM.

**Fig S6.**
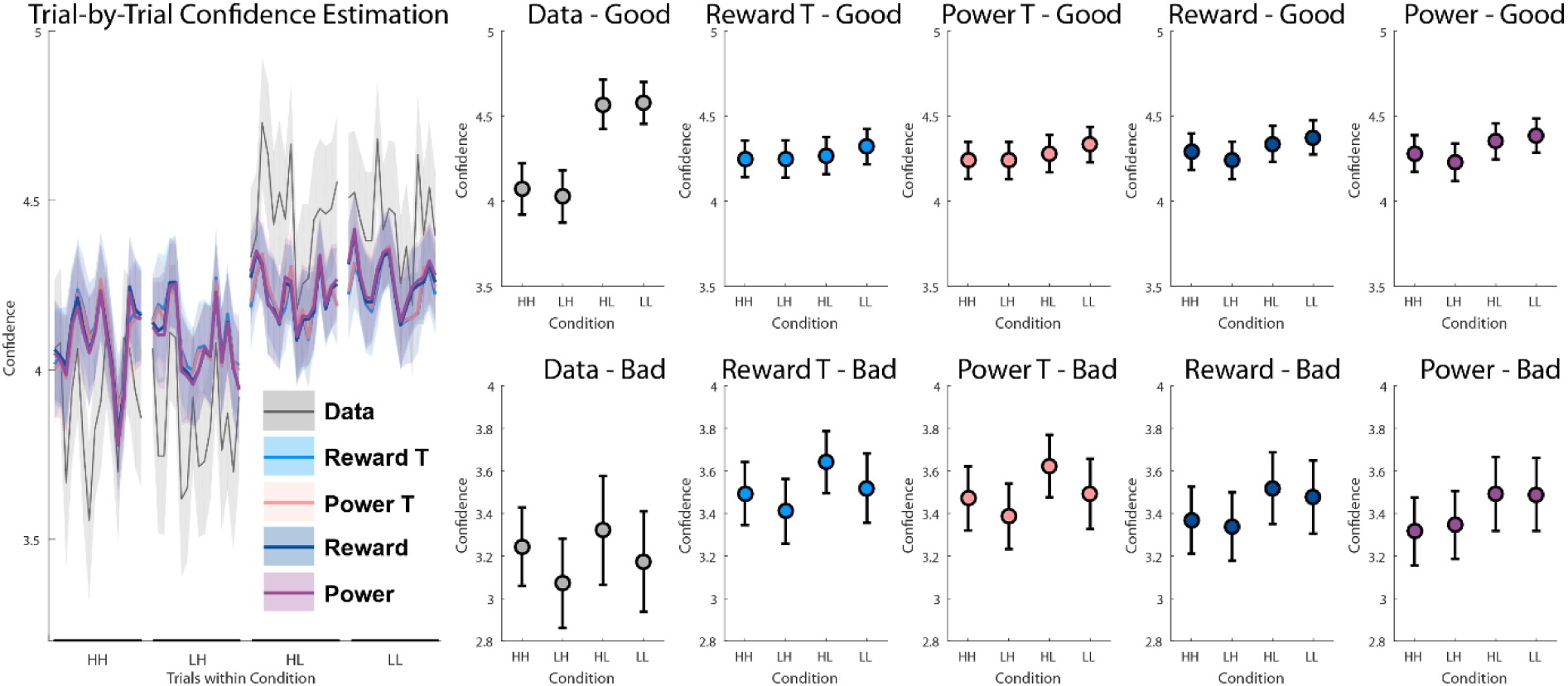
Power Utility Model Performance in Experiment 1.

To examine other utility functions we fitted a utility model which transforms the rewards in each trial according to the power utility function [1,2]:

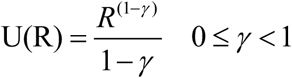

Where *γ* is the risk aversion factor – the closer it is to 1 the participant is more risk averse (and the closer the function is to log(r)). We used a model that learns from these transformed values, i.e. from utilities and not directly from the rewards, but was otherwise exactly the same as the ‘Reward’ model:

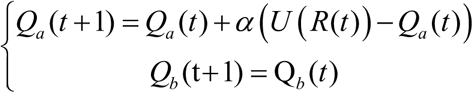

When option a is chosen, its value is updated according to the difference between its current value and the utility of the option’s current reward, with a learning rate *α*. Decision in each trial was then carried using a softmax rule.

We fitted this ‘Power’ model to the choice data, and an additional ‘Power-T’ model which added the drift to threshold of the unchosen option mechanism:

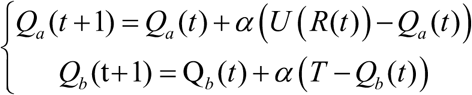

We examined the fit of these models to the data and how well they predicted the confidence reports.

We found that the ‘Power’ and ‘Power-T’ models performed very similarly to the ‘Reward’ and ‘Reward-T’ models respectively. Their WAIC values were: ‘Power’ 228.06 ± 67.46, ‘Power-T’: 214.51 ± 68.66, whereas the ‘Reward’ models WAIC values were: ‘Reward’ 226.55 ± 67.67, ‘Reward-T’ 214.57 ± 68.79. ‘Power-T’ was as good as our best model in explaining choice behaviour.

We than examined how well the ‘Power’ models explained confidence reports. Again, they fared similarly to the ‘Reward’ models with linear fit (R2) of: Power’ 0.21 ± 0.22, ‘Power-T’: 0.21 ± 0.21, whereas the ‘Reward’ models WAIC values were: ‘Reward’ 0.21 ± 0.22, ‘Reward-T’ 0.21 ± 0.21.

Finally, we examined the predicted confidence reports in the four condition blocks, and found that the patterns predicted by the ‘Power-T’ model were identical to the pattern predicted by the ‘Reward’ model. We concluded that the transformation of reward in a trial by trial manner did not introduce any new mechanism to learning beyond the one already implemented by the ‘Reward’ model.

**Fig S7.**
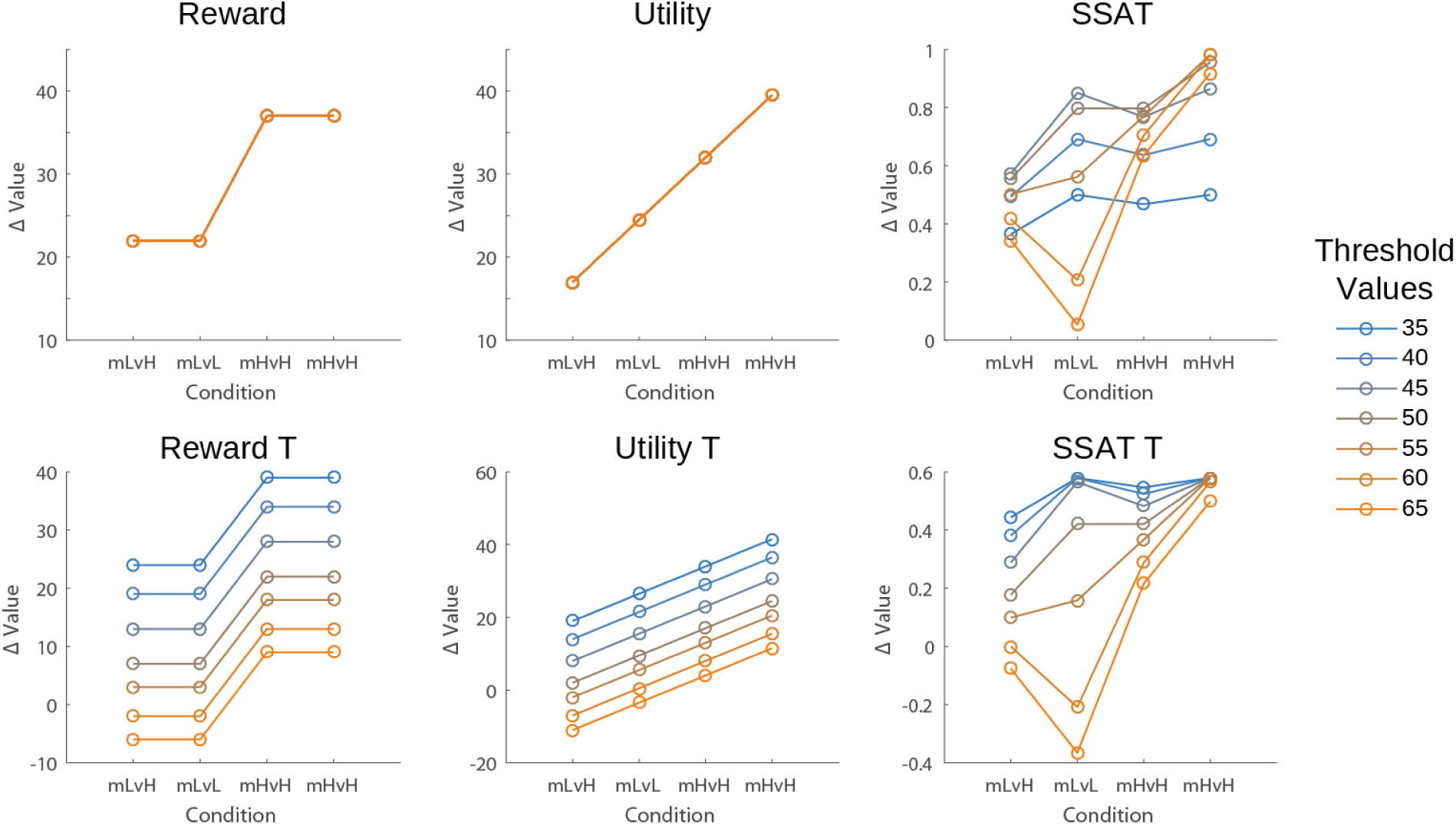
Value-Differences Optimality Analysis in Experiment 2. We followed the same optimality analysis as in Figure S3 with the reward distributions from experiment 2, and varied the acceptability threshold (blue to yellow lines). We found that the models varied dramatically in the relative values they assigned the options. Again, a greedy decision maker would therefore be able to accumulate similar amount of rewards using each model. However, a noisy decision maker (modeled using softmax) may be more likely to choose the low mean reward option in some conditions, according to the models’ predictions.

**Fig S8.**
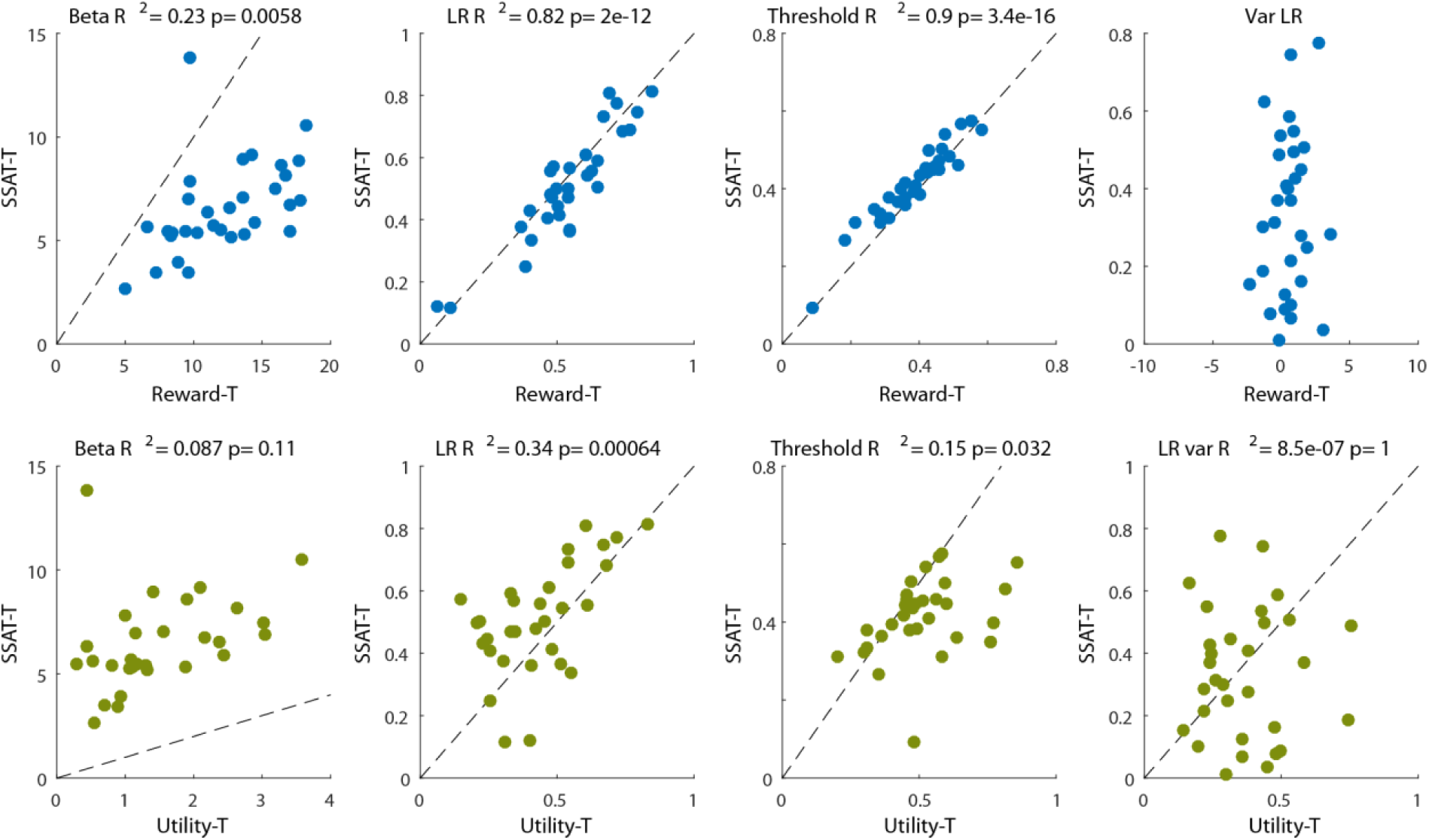
Relations between estimated parameters in different models in experiment 2. We compared the individual parameters estimated for each of our drift (and best performing) models. The Reward-T and SSAT-T models’ parameters for learning rates and threshold were almost identical for all our participants. Reward model gave the best fit to decisions, while the SSAT-T model gave the best fit to confidence reports. This indicates that these models may use a shared mechanism for decisions, but the SSAT-T model uses the reward variance information to generate confidence reports. Parameters estimations were not as similar for the SSAT-T and the Utility-T.

**Fig S9.**
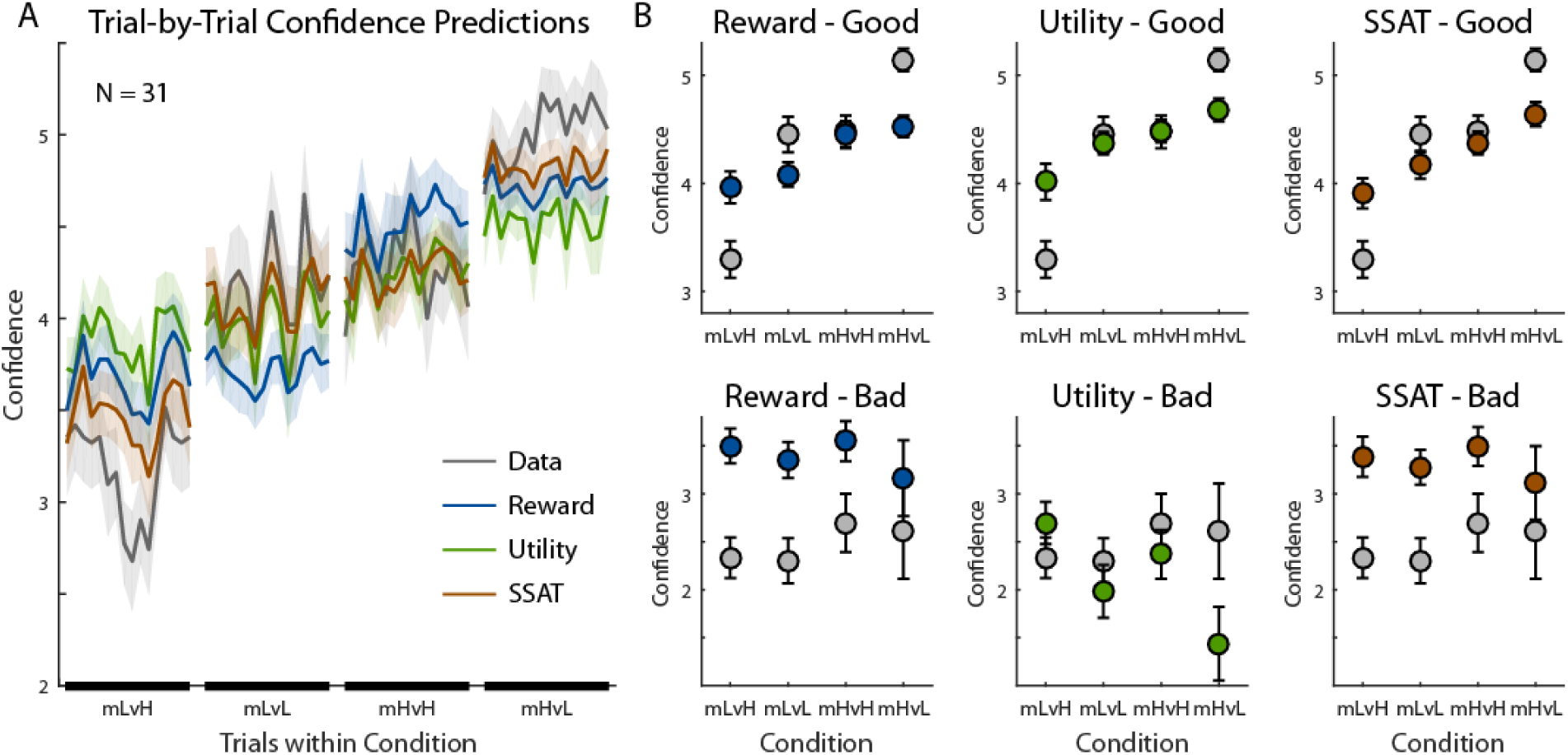
Model predictions for confidence reports in experiment 2. (A) Trial-by-Trial confidence reports (grey line) and model predictions during each experimental condition are displayed, averaged across participants (shaded areas represent SEM). (B) Models’ predictions for confidence reports when choosing the good option (Top Row) and when choosing the bad option (bottom row). Predictions were averaged between trials 10-25 in each block. The average reports made by participants is displayed in grey. All models predicted higher confidence when choosing the good option than when choosing the bad option. Error bars represent SEM.

